# Roles of Clamp Closing and Allosteric Effects of Discriminator and Upstream Interactions on Downstream Elements in Stabilizing an *E. coli* RNA Polymerase-Promoter Open Complex

**DOI:** 10.1101/2022.04.18.488589

**Authors:** Hao-Che Wang, Krysta Stronek, M. Thomas Record

## Abstract

*E. coli* RNA polymerase (α_2_ββ’ωσ^70^) forms stable open complexes (OC) at λP_R_ and T7A1 promoters at 37 °C at similar rates but with very different lifetimes. To probe the origins of these differences, we determine OC lifetimes for full-length (FL) and downstream-truncated (DT) λP_R_ and T7A1 promoter variants with eight combinations of discriminators and upstream elements. We find the discriminator is the major determinant of OC lifetime, while upstream elements modulate the discriminator effect. The very different lifetimes of these stable OC arise primarily from differences in 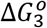, the free energy change for converting I_2_ (the initial open intermediate) to stable OC. 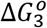 is twice as favorable for λP_R_-discriminator variants as for T7A1-discriminator variants. Truncation at +6 (DT+6) eliminates most differences in 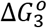, destabilizing λP_R_-discriminator variants without affecting T7A1-discriminator variants. All DT+6 OC are much more stable and longer-lived than I_2_. Urea greatly destabilizes OC, primarily affecting RNAP interactions with +6 and upstream DNA. From these findings we deduce that stabilization of I_2_ involves closing the β’-clamp with extensive coupled folding to contact the region of promoter DNA from the transcription start-site to +6. Lifetime differences between DT+6, DT+12 and FL promoters allow dissection of contributions to OC stability from interactions of regions of the downstream duplex with downstream mobile elements (DME; β-lobe, β’–clamp, β’-jaw). We propose an allosteric network by which differences in discriminator and −10 sequence are sensed by σ_1.2_ and β-gate loop and transmitted to σ_1.1_ and β-lobe to affect DME-duplex interactions that determine OC lifetime.

## Introduction

Regulation of transcription occurs at all stages from initiation to termination. The first and most fundamental level of regulation is promoter sequence. *In vivo*, initiation rates vary at least 10,000-fold for different promoters, from ~1 s^-1^ to once per generation or less. Rates of open complex (OC) formation (compared at the same RNA polymerase (RNAP) concentration) and dissociation *in vitro* span similar ranges determined by promoter DNA sequence [1–4]. Transcription in *Escherichia coli* begins with formation of an initial specific (closed) complex between RNAP holoenzyme (α_2_ββ’ωσ^70^) and duplex promoter DNA. At the *λ*P_R_ promoter, as summarized in Fig. 1 (see [5] and [6]), kinetic-mechanistic evidence indicates that initial binding triggers a series of conformational changes beginning with bending and wrapping of far-upstream duplex DNA (the UP element and beyond) on RNAP to contact the β’ clamp and form an ensemble of intermediate closed complexes collectively designated as I_1_. In the most advanced I_1_ species, the downstream promoter duplex is bent into the RNAP cleft and the β’ clamp is closed. In the subsequent rate-determining step, the clamp and 13 bp of promoter DNA including the transcription start site (TSS) open together to form the initial short-lived open intermediate I_2_. Other mechanisms of OC formation have been proposed based on kinetic studies with T7A1 [7–9] and other promoters [10–13], and from cryoEM studies of RNAP-promoter complexes with [14] and without protein factors [15, 16]. At *λ*P_R_ and T7A1 promoters, additional conformational changes convert I_2_ to longer-lived, more stable OC (for *λ*P_R_ called I_3_ and RP_O_) [3, 17–20]. For *λ*P_R_, RP_O_ (the stable 37 °C OC) cannot initiate, and must convert to the I_3_ intermediate in order to initiate [21, 22].

**Figure 1.**
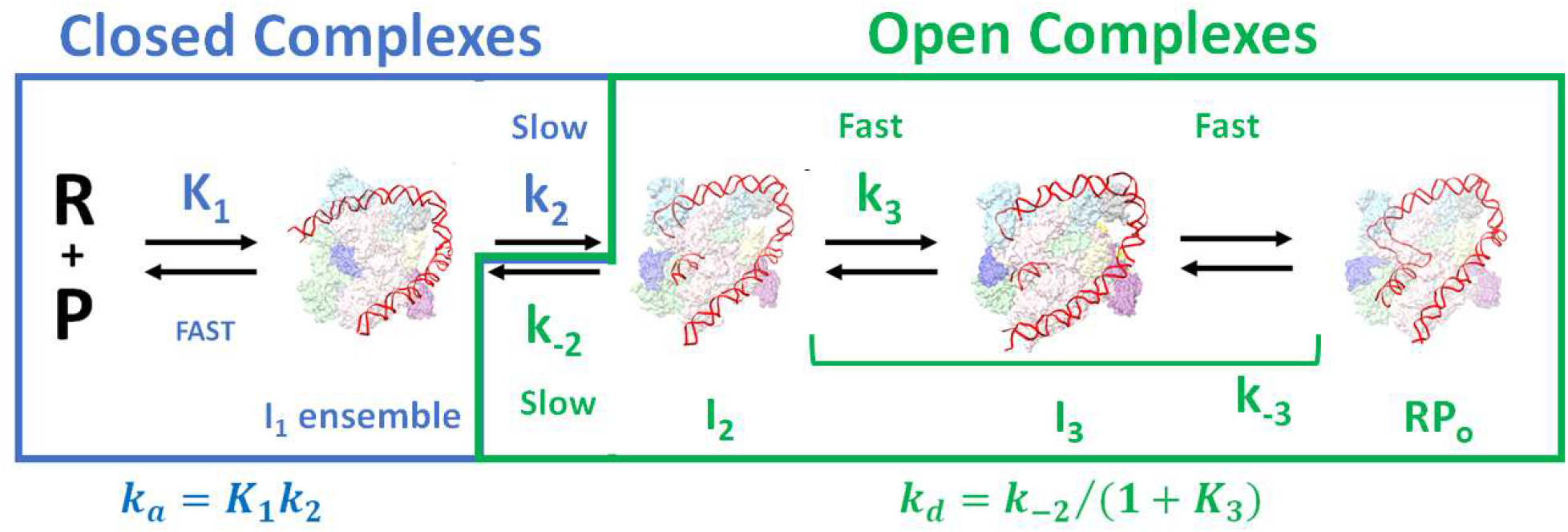
Intermediates in Formation and Dissociation of the 37 °C Stable RP_O_ Complex between RNAP and λP_R_ Promoter DNA. Significant open-promoter intermediates (I_3_, I_2_) identified in the mechanism of dissociation of the 37 °C open complex (RP_O_) and their contribution to the dissociation rate constant k_d_ = k_-2_/(1+K_3_) and RP_O_ lifetime 1/k_d_ are shown in green. Significant closed-intermediates (RP_C_ and other closed complexes in the I_1_ ensemble) identified in the mechanism of open complex formation and their contribution to the overall 2^nd^ order association rate constant k_a_ = K_1_k_2_ are shown in blue. The DNA opening-DNA closing step in midmechanism, like the catalytic step of an enzyme-catalyzed reaction, is rate-determining in both directions because the preceding steps (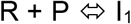 ensemble; 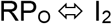)) are rapidly reversible on the time scale of the relevant direction of this step. (I_1_ ensemble is generated from PDB 6PSQ [14], I_2_ and I_3_ are generated from PDB 6PSV [14] and RP_o_ is generated from PDB 7MKD [22]. The wrapping DNA and σ_1.1_ region (PDB:1SIG [62]) are incorporated to the indicated PDB models by PyMOL ver. 2.3.2 and are illustrated by CSF ChimeraX version 0.91.)

The goal of the research presented here is to discover the interactions and conformational changes responsible for the very different lifetimes of stable λP_R_ and T7A1 OC from one another and from I_2_ intermediates, and thereby understand the process of conversion of I_2_ to stable OC. To accomplish this, we determine lifetimes of stable OC formed with full-length (FL) and downstream-truncated (DT) *λ*P_R_ and T7A1 promoter constructs with interchanged discriminators and upstream elements, as well as I_2_ lifetimes for FL promoters, to complement previous determinations with wild-type *λ*P_R_ and T7A1 constructs [3, 17, 20].

From fast footprinting and filter binding (FB) kinetic studies on the *λ*P_R_ promoter, we previously deduced that conversion of I_2_ to I_3_ and RP_O_ involved repositioning the discriminator non-template strand in the cleft and assembly of RNAP downstream mobile elements (DME) on the initial transcribed region (ITR, +3 to +20) of the promoter duplex [3, 18–20, 23]. Deletion of the β’ jaw domain DME significantly reduced the lifetime of the 37 °C *λ*P_R_ OC (RP_O_) [3, 20], but had only a small effect on the lifetime of the less stable 37 °C T7A1 OC [3]. Two in-cleft RNAP mobile elements, both essential for growth [24, 25], also have large effects on lifetime. Deletion of the β gate loop (β GL), which is positioned across from the β’ clamp in the DNA channel, resulted in a 22-fold reduction in the lifetime of the stable *λ*P_R_ OC, but only a 4-fold lifetime reduction for the T7A1 OC [26]. Conversely, deletion of σ_1.1_, a RNAP mobile element that is ejected from the downstream helix channel during the formation of a stable OC at *lacCONS* [27] and *λ*P_R_ (but not T7A1 [3]) promoters, greatly stabilizes the T7A1 OC but has little effect on stability of the already-very-stable *λ*P_R_ OC [3].

Truncation of the *λ*P_R_ promoter at +12 reduced the lifetime of the 37 °C OC (RP_O_) by a similar amount to that observed for ΔJAW RNAP [20], but had only a small effect on the lifetime of the less stable 37 °C T7A1 OC [3, 20]. For the N25cons promoter, single base pair truncations of the downstream helix from +21 to +6 progressively reduced OC lifetime [28]. Further truncation upstream of +6 increased OC lifetime but resulted in a complex that was incapable of initiation upon NTP addition [28].

The discriminator, a 4-8 base promoter element that separates the −10 element from the transcription start site (TSS), is also a major determinant of OC lifetime [29–31]. OC lifetimes of hybrid promoters made by exchanging λP_R_ and T7A1 discriminators are similar to those of the parent promoter constructs with the same discriminator [31]. Recent cryoEM structures of stable *λ*P_R_ OC show that a 6 base discriminator like that of *λ*P_R_ bridges the gap between the −10 region and the transcription start site without distortion [22], and a 6 base discriminator is the most common discriminator length [32, 33]. Longer discriminators (like T7A1; 7 bases) require pre-scrunching and shorter discriminators require stretching to put the start-side base in the active site [15, 34–36]. The intrinsic effect of pre-scrunching on OC lifetime has not been determined, but pre-scrunching of 7 base discriminator strands was deduced to reduce OC stability by about 0.7 kcal [34].

The base sequence near the upstream end of the discriminator has a major effect on OC lifetime. For WT and variant versions of five promoters (including λP_R_ and *rrnB* P1), OC lifetime is 7-50 fold greater with a non-template strand G or A than with C at the 2^nd^ position from the upstream end [29, 30]. Comparison of A and G variants at this position for two promoters reveal only small lifetime differences (0.5-1.5-fold). Alanine mutational scanning shows that this stabilization results from the interaction of the 2^nd^ position G base with σ_1.2_ residues Y101 and M102 [30]. Base identity at the 3^rd^ and 4^th^ positions of the discriminator also significantly affects OC lifetime. For stable OC formed by WT RNAP at the *λ*P_R_ promoter, changing both the 3^rd^ and 4^th^ non-template strand bases from T to C reduces lifetime 4-fold, while replacement of either T by A reduces lifetime by 20-30% [26]. In all cases, lifetime differences between WT and variant promoters were minimized (< 2-fold) when investigated with the Δ gate loop (GL) RNAP variant, indicating that the β GL interacts with these positions of the discriminator, and MnO_4_^-^ reactivities of thymines at −3 and −4 positions of the *λ*P_R_ discriminator were significantly higher in OC with WT RNAP than with ΔGL RNAP [26].

Building on previous research on the determinants of open complex lifetime and stability for λP_R_ and T7A1 promoters, here we investigate a set of hybrid promoter constructs with different combinations of the λP_R_ and T7A1 discriminators, core regions and UP-elements. We obtain dissociation rate constants for both initial OC (I_2_) and stable OC (e.g. λP_R_ I_3_, RP_O_) formed by wildtype (WT) RNAP at these promoters. From analysis of these dissociation rate constants, we obtain equilibrium constants K_3_ for the conversion I_2_ → stable OC at each promoter and the corresponding standard free energy change 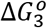 for this process. We also report dissociation rate constants for stable OC formed by WT RNAP with downstream truncations of these promoters at +6 and +12, and for stable OC formed at FL promoters by a deletion mutant RNAP lacking the downstream mobile jaw. Analysis of K_3_ for downstream-truncated promoters yields contributions to 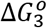 from interactions of downstream mobile elements (DME: downstream β’ clamp, β lobe, β’ jaw) of RNAP with three regions of the downstream duplex (+6 and upstream, +7 to +12, downstream of +12). Differences in these free energy contributions for different discriminators and upstream elements provide insight into the allosteric network by which upstream sequence information is communicated to these DME to determine OC lifetime. Consistent with previous research [3, 26, 29, 30], we deduce that interactions of σ_1.2_ and the β GL with non-template strand bases of the upstream discriminator are the key upstream interactions in this allosteric mechanism. These upstream interactions of σ_1.2_ and the β GL with specific discriminator bases are transmitted downstream to affect DME interactions with the promoter duplex downstream of +6. We also determine and dissect effects of the destabilizer urea on 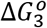 for T7A1 and λP_R_ promoters and interpret these results in terms of coupled folding of regions of the β’ clamp when it closes on the downstream duplex (+3 to +6) and transcription start site (TSS) in the conversion of I_2_ open intermediates at these promoters to stable OC.

## Results

### Lifetimes of Open Complexes at λP_R_ and T7A1 Promoter Variants

To characterize the interactions responsible for lifetimes and stabilities of open complexes, and the roles of different regions of the promoter as determinants of OC lifetime and stability, we studied a family of eight λP_R_ and T7A1 promoter variants with different combinations of discriminators, core promoters (including −10 and −35 elements) and UP-elements. Each region (UP-element, core promoter, discriminator, in that order) is designated as L (λP_R_) or T (T7A1), so TTL is the promoter variant with the UP-element and core promoter element of T7A1 and the λP_R_ discriminator. Parent λP_R_ (LLL) and T7A1 (TTT) promoter sequences and the division into three regions are shown in Fig. 2A. In all cases the downstream region (initial transcribed region, ITR) is a modified version of the λP_R_ ITR, used previously in related experiments [21, 31, 37].

**Figure 2.**
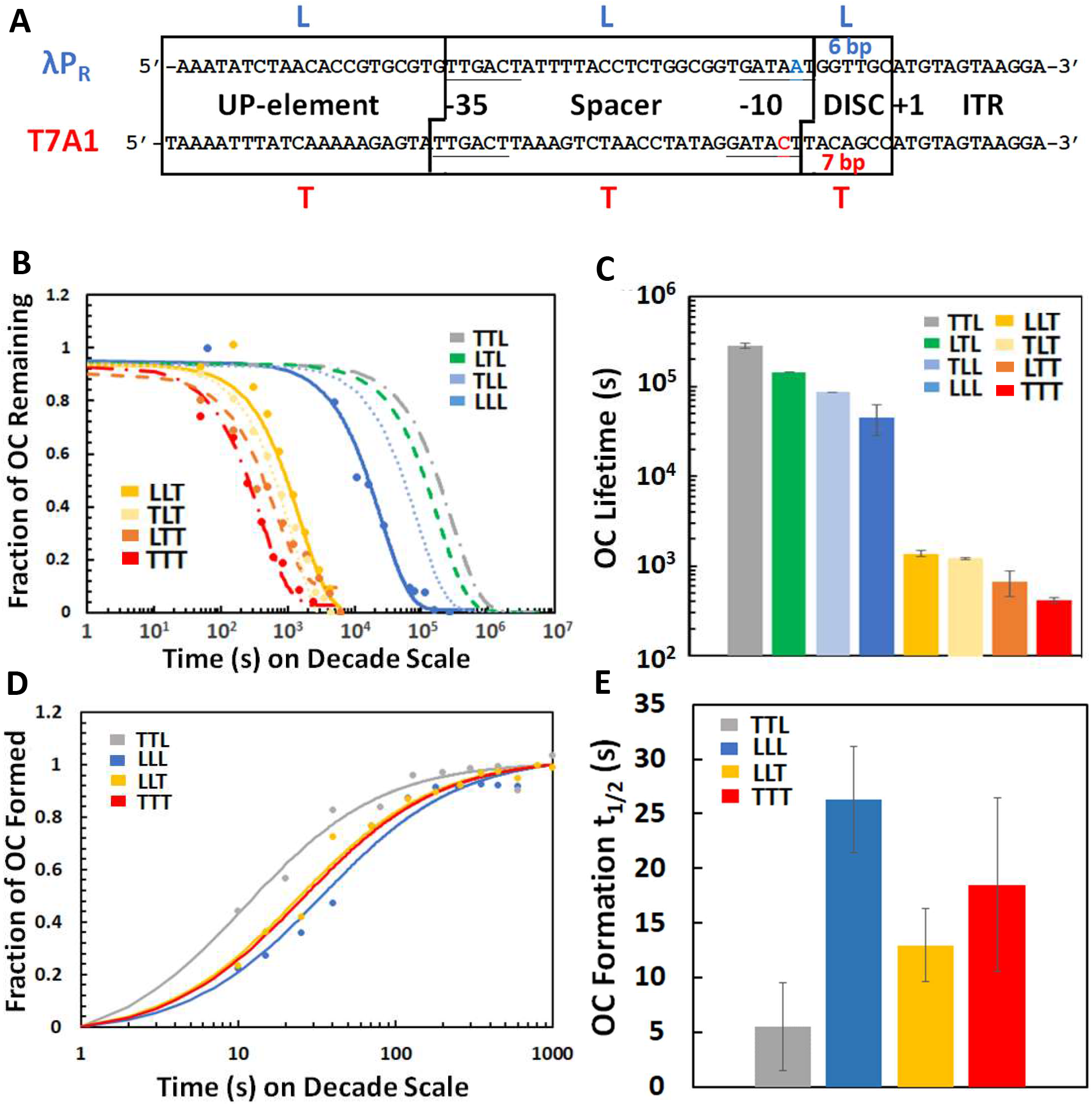
OC Lifetimes and Halftimes of OC Formation of λP_R_ and T7A1 Promoter Variants. A) Non-template strand sequences of central λP_R_ (abbreviated LLL) or T7A1 (abbreviated TTT) promoters embedded in nonspecific DNA fragments (λP_R_: −80 to +42, T7A1: −81 to +42). Discriminator, core region (including −10, spacer and −35 elements), and UP-element are indicated; variant promoters with all combinations of these three regions are investigated here. The promoter with three λP_R_ regions is designated LLL and that with three T7A1 regions is designated TTT. The initial transcribed region (ITR) of both promoters is modified from that of λP_R_ [31] B) Fraction of OC remaining vs time (log scale) from filter binding dissociation kinetic assays. Curves for TTL, LTL and TLL are obtained by extrapolation of dissociation kinetic data as a function of [urea], as described subsequently. C) Log scale histogram of OC lifetimes for hybrid promoter constructs. Promoters are ranked in an analogous sequence order in both discriminator series, with LLL and TTT at the right end and TTL and LLT at the left end of each group. D) Fraction of OC formed at TTL, LLL, LLT and TTT promoters vs. time (log scale), determined from filter binding association kinetic assays performed at 10 nM RNAP and 10 nM promoter DNA. E) Histogram of half times t_1/2_ of OC formation (t_1/2_ = 1/(k_a_[RNAP]) for TTL, LLL, LLT and TTT promoters from results of Fig 2D.

These promoters differ in multiple ways, from their discriminators to their UP-elements, as shown in Fig. 2A. Discriminators differ in length by 1 bp (6 bp L; 7 bp T) as well as in sequence. At the upstream end, the L discriminator begins with GG and the T discriminator begins with TA. The core L and T promoters are quite similar, with the same −35 sequences and the same spacer length (17 bp). The −10 sequences differ at the position adjacent to the downstream end (position −8 for a 6 bp discriminator, as in λP_R_). At this position, λP_R_ has A, the consensus base, and T7A1 has C. Both halves of the T UP-element are similar to consensus, being AT-rich with A tracts, while only the distal half of the L UP-element is AT-rich, with a shorter A tract.

Fig. 2B shows time courses (on a log time scale) of irreversible OC dissociation for these 8 promoters, determined by the nitrocellulose filter binding assay described in Methods. Kinetics are first-order, yielding dissociation rate constants k_d_ and OC lifetimes (τ_d_ =1/k_d_). Values of k_d_ are listed in Table S1. Curves plotted in Fig. 2B correspond to these τ_d_ values. Exceptionally long-lived OC for promoters with the L discriminator were studied at different urea concentrations, as described below, and the results extrapolated to zero urea concentration to obtain τ_d_ and predict the kinetic curves shown for these variants in Fig. 2B. Lifetimes τ_d_ of stable OC are compared in the log-scale bar graph in Fig. 2C.

Lifetimes of these promoter OC cluster into two groups determined by their discriminator element (Fig. 2C; Table S1). The lifetime of LLL exceeds that of TTT by somewhat more than two orders of magnitude (~130-fold). The effect of changing from T-discriminator to L-discriminator in a variant promoter, retaining the same core and upstream elements, can be either much greater or much less than 130-fold. The change from TTT to TTL has the largest effect on τ_d_ (690-fold), while the change from LLT to LLL has the least effect (40-fold). Changes from LTT to LTL and from TLT to TLL have intermediate effects (290-fold, 70-fold respectively). These highly variable effects of changing discriminator follow from the unexpected observation that within each set of four promoters with the same discriminator, the OC formed by the parent promoter (TTT, LLL) has the smallest τ_d_. For both L- and T-discriminator series, the order of τ_d_ is the same: WT < UP-element change < core-element change < UP- and core element change (i.e. TTT < LTT < TLT < LLT and LLL < TLL < LTL < TTL). Lifetimes of stable OC at these L- and T-discriminator promoters span ~5-fold and ~3-fold ranges, respectively.

For the L-discriminator series, changing the core promoter from L to T results in a moderate (~3.5-fold) increase in τ_d_; this increase is the same whether the adjacent UP element is T or L (see Fig. S1A). The increase in τ_d_ for changing the UP element from L to T in the L-discriminator series is more modest (~1.5-fold), and also is the same in the context of either T or L core promoter. For the T-discriminator series, effects of changes in UP element and core promoter are mostly smaller than for the L series and do not appear independent (see Fig. S1B). Changing the core promoter from T to L increases τ_d_ slightly more starting with TTT(~3-fold) than starting with LTT (~2-fold). Changing the UP-element of the TTT promoter to L increases τ_d_ by ~50%, but changing the UP-element of the TLT promoter to L increases τ_d_ minimally (< 20%).

### Half-Times for Formation and Stabilities of Open Complexes at λP_R_ and T7A1 Promoter Variants

Time courses of determined by filter binding as described in Methods, are shown in Fig. 2D. Data were fit as described in Methods to obtain composite 2^nd^ order association rate constants k_a_ (defined in Fig 1). Half-times t_1/2_ of OC formation at these equimolar reactant concentrations are calculated from k_a_ (see Table S1) and compared on the bar graph of Fig. 2E. The t_1/2_ for OC formation at LLL is slightly (~1.5-fold) larger than for TTT. This small difference in t_1/2_ is much less than the difference in τ_d_ (~130-fold) between LLL and TTT OC. For LLL and, to a lesser extent, TTT promoters, swapping in the core and UP elements of the other promoter affects t_1/2_ moderately. The association t_1/2_ of LLL is ~5-fold greater than TTL, while that of TTT is ~2/3 that of LTT.

As a result of the large differences in OC lifetime (τ_d_) and smaller (opposite direction) differences in t_1/2_ for OC formation between promoters with T- vs. L-discriminators and the same upstream elements, the overall equilibrium constant for stable OC formation from RNAP and promoter DNA (K_1_K_2_K_3_ = k_a_/k_d_; see Fig. 1) is ~20-fold larger for LLL than for LLT, and ~2000-fold larger for TTL than for TTT (Table S1). Comparing the effect of changing upstream elements while keeping the discriminator the same, K_1_K_2_K_3_ is ~25 times larger for TTL than for LLL, and ~5 times larger for LLT than for TTT (see Table S1).

### Dissociation Lifetimes (Bubble Closing Rates) of Open Intermediates I_2_

In principle, the large differences in OC lifetime of the different variant promoters with T and L discriminators could arise from different rates of bubble closing (I_2_ → I_1_; rate constant k_-2_) and/or different equilibrium constants K_3_ for OC stabilization (I_2_ → stable OC; see Fig. 1). From a molecular perspective, differences in strength and/or extent of downstream interactions with the duplex and/or from differences in upstream interactions with the discriminator and −10 regions could be responsible for the very different lifetimes of stable OC formed with these promoter variants. To separate these contributions, we determined DNA bubble-closing rate constants (k_-2_) after rapid KCl-upshifts that destabilize the OC. The KCl-upshift creates a large transient population (burst) of the intermediate OC (I_2_) which subsequently decays in the irreversible bubble-closing step [5, 17]. Filter-binding measurements on samples quenched as a function of time after the salt upshift (see Methods) allow determination of the rate of bubble-closing and the lifetime of I_2_, previously found to be independent of KCl concentration [17]. Fig. 3A shows that decays of I_2_ are single-exponential, yielding I_2_ lifetimes (τ_-2_ = 1/k_-2_; see Figure 3B and Table S1) which span a 10-fold range. For three promoters with the T discriminator (TTT, LTT, LLT) and two promoters with the L-discriminator (TTL, TLL), τ_-2_ is in the range 1 to 2 s. Bubble closing is marginally slower for TLT (τ_-2_ = 5 ± 3 s) and faster for LTL and LLL (τ_-2_ = 0.4 ± 0.3 s). Differences in τ_-2_ between L- and T-discriminator variants with the same upstream elements appear significant in some cases but are much smaller than (and in the opposite direction from) differences in τ_d_ for dissociation of stable OC formed by these L- and T-discriminator variants.

**Figure 3.**
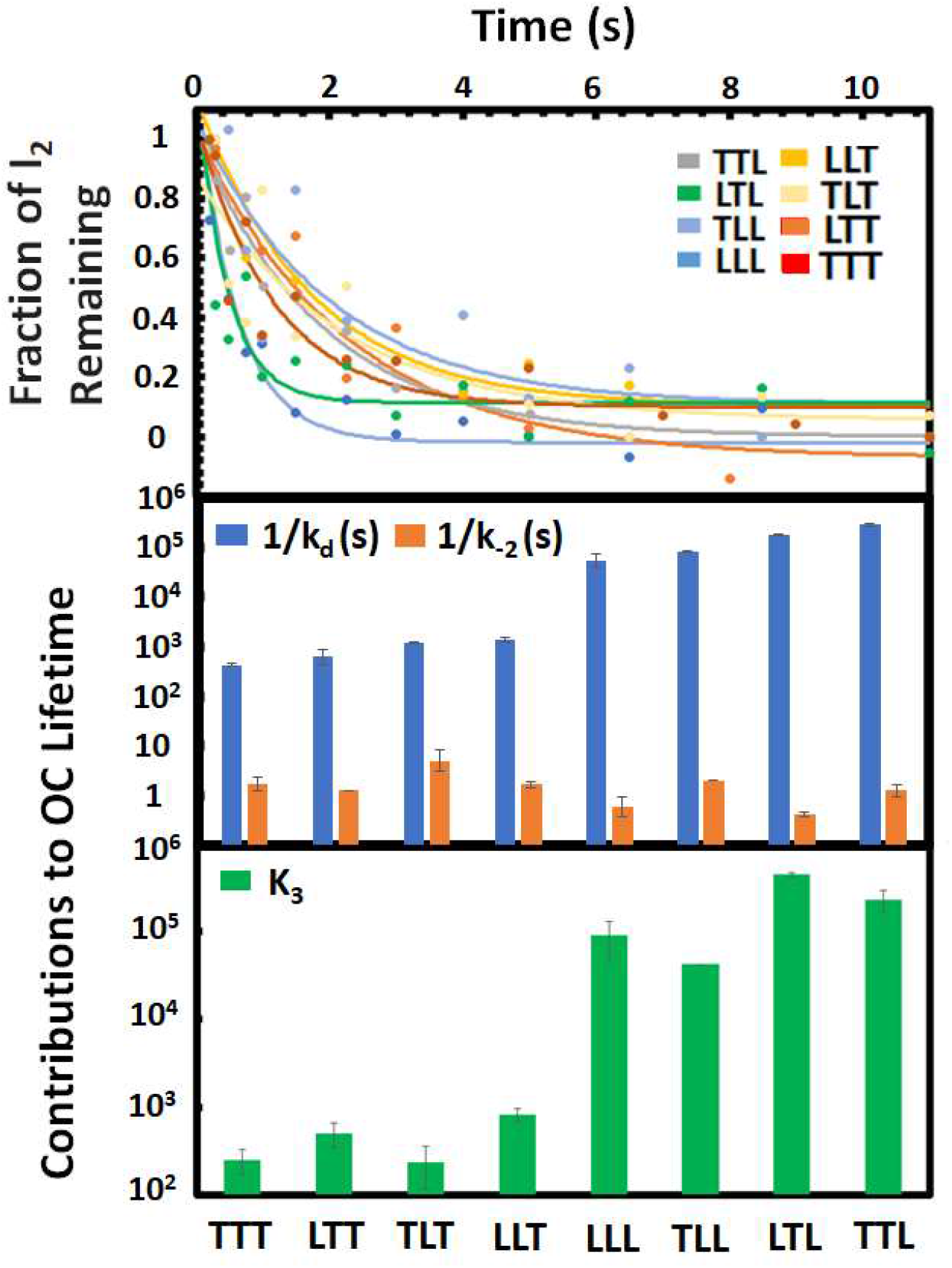
Contributions to Lifetime of the Stable OC. A) Filter-binding dissociation kinetic assays after rapid OC destabilization by high [salt] to determine dissociation lifetimes of OC intermediates I_2_ formed with variant promoters [17]. B) Histograms comparing dissociation lifetimes (s, on decade scale) of the stable OC (1/k_d_, blue) and of the intermediate I_2_ (1/k_-2_, orange). C) Equilibrium constants K_3_ ≅ k_-2_/k_d_ (on decade scale) for conversion of I_2_ to the stable OC.

Equilibrium constants K_1_K_2_ = k_a_/k_-2_ (Fig. 1) for the conversion of free RNAP and promoter to the I_2_ intermediate OC at each promoter are listed in Table S1. Propagated uncertainties in these values are large (50-80%) but it is clear that K_1_K_2_ values for forming the I_2_ intermediate span at least one order of magnitude, in the order TTL > LLT > TTT > LLL. As observed for stability and lifetime of the stable OC, stabilities of I_2_ complexes formed with the TTL and LLT variants exceed those of the parent TTT and LLL promoters.

### Equilibrium Constant K_3_ and Free Energy Changes 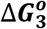 for Conversion of I_2_ Open Intermediate to a Stable OC

From the lifetimes of stable OC (τ_d_ = 1/k_d_) and of the I_2_ intermediate (τ_-2_ = 1/k_-2_) for the sets of FL and DT promoters with L and T discriminators investigated here, values of equilibrium constants K_3_ (defined in Fig. 1) and corresponding standard free energy changes 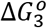 for the conversion I_2_ → stable OC are readily determined. Values of K_3_ (Table S1) for promoters with the L-discriminator span a 4-fold range and are 100 to 1000-fold larger than for promoters with the T-discriminator, which also span a 4-fold range. As observed for k_d_ above, the discriminator is the most important determinant of K_3_ (100- to 1000-fold effect on K_3_ of replacing T by L discriminator), followed distantly by the choice of core promoter (1 to 5-fold), while the choice of UP element has relatively little effect (< 1.5-fold) on K_3_.

### Assessing Contributions of Interactions with Different Downstream Regions to Lifetimes (1/k_d_) of Open Complexes for Variants with T or L Discriminator

To determine contributions from interactions of RNAP elements with different downstream regions of these promoter variants, we determined effects on OC lifetime of truncating the downstream duplex at +12 or +6, or of deletion of the β’ downstream mobile jaw (ΔJAW). These results (Fig. S2) are summarized in the bar graphs of Fig. 4 and in Table S2.

**Figure 4.**
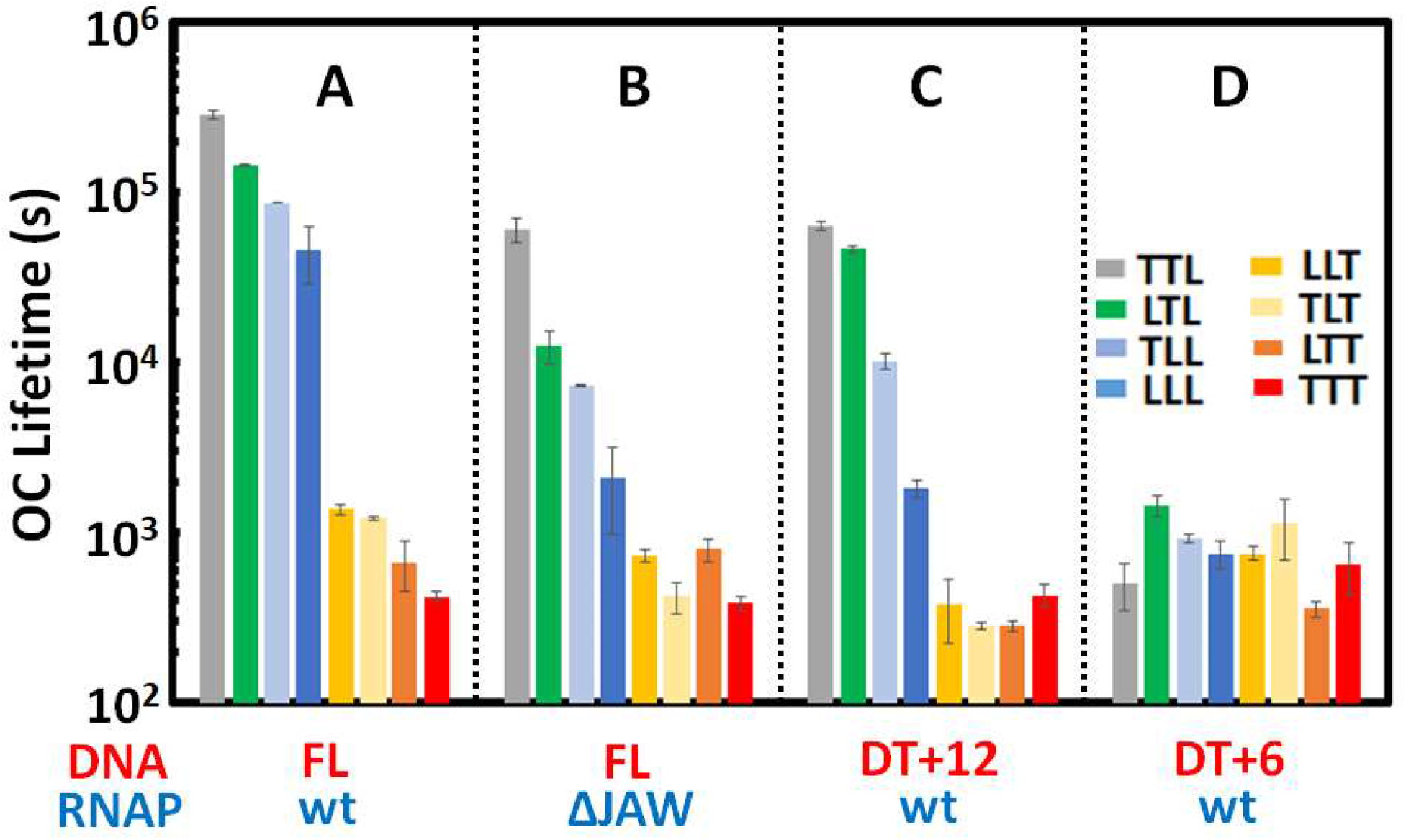
Comparisons of OC Lifetimes for WT RNAP and Downstream-Truncated (DT) Promoter Variants and for ΔJAW RNAP and Full-length Promoter Variants. Decade-scale histograms compare OC lifetimes (in seconds) for A) intact RNAP and full-length (FL) λP_R_ promoter DNA; B) ΔJAW RNAP and FL λP_R_ promoter DNA; C) intact RNAP and DT+12 λP_R_ promoter DNA and D) intact RNAP and DT+6 λP_R_ promoter DNA.

Most notably, while downstream truncation of the promoter (to +12 or +6) or deletion of the downstream mobile jaw (ΔJAW) greatly reduces lifetime of stable OC formed at L-discriminator series promoters, these deletions/truncations have little effect on the lifetime of stable OC formed at T-discriminator series promoters. Analogous results for LLL (DT+12, ΔJAW) and TTT (ΔJAW) promoters were reported previously [3, 20]. For L-discriminator series promoters, Fig 4 B and C show that deletion of the downstream jaw and truncation of the promoter DNA at +12 have similarly large effects, reducing OC lifetime by 0.5 to 1.5 orders of magnitude. The smallest effects are on TTL and the largest on LLL. Additional large reductions in lifetime are observed by truncation of L-discriminator series promoters at +6 (Fig. 4D), while truncation of T-discriminator series promoters at +6 has little effect on OC lifetime.

Fig. 4D (see also Table S2) shows that lifetimes of DT+6 OC are similar for all promoter variants investigated. Comparison with results for the I_2_ intermediate OC at these promoters in Figure 3B and Table S1 reveals that in all cases lifetimes of DT+6 OC greatly exceed those of I_2_. (Table S1 and S2). This comparison indicates that contacts with the region of the promoter upstream of +6 greatly stabilize the OC, as demonstrated below.

### Contributions of RNAP Interactions with Different Downstream Regions to the Equilibrium Constant K_3_ and Standard Free Energy Change 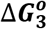 for OC Stabilization (I_2_ → stable OC)

For FL promoter variants and WT RNAP, equilibrium constants K_3_ for conversion of the I_2_ open intermediate to a stable OC (obtained from K_3_ ≅ k_-2_/k_d_; Fig 1) were reported in Fig 3C. Corresponding values of K_3_ for DT+12 and DT+6 variants of these promoters, calculated from their k_d_ values (Table S2) and their DNA closing rate constants k_-2_ (Table S1), are reported in Table S3. Standard free energy changes 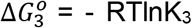 for formation of a stable OC on FL promoter DNA from the open I_2_ intermediate and contributions to 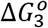 from interactions of RNAP with three downstream regions of the promoter (downstream of +12; +12 to +7; upstream of +6), calculated from these K_3_ values for all eight FL and their DT+12 and DT+6 variants at 37 C, are reported in the bar graphs of Fig. 5 (see also Table S4).

**Figure 5.**
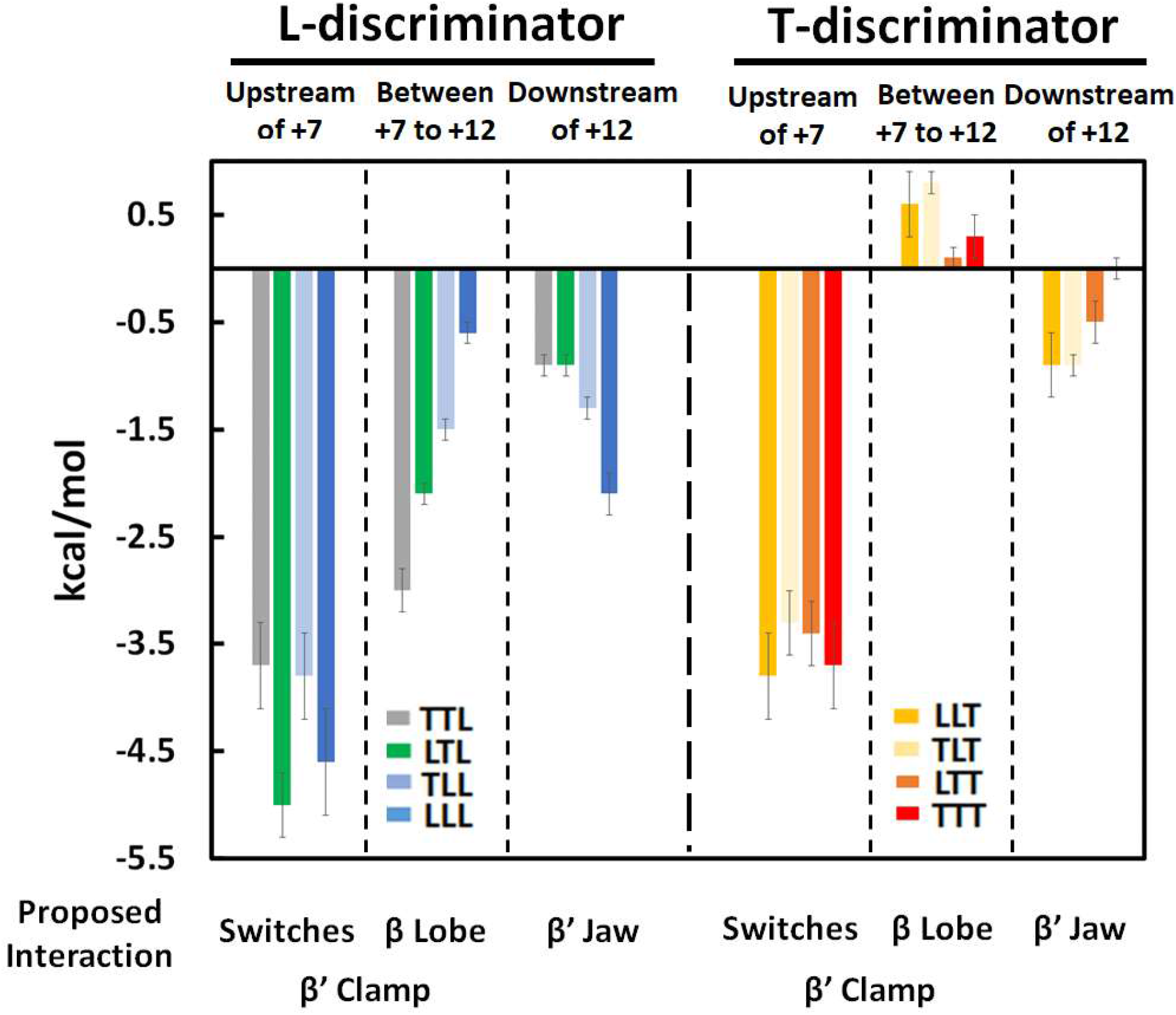
Contributions to the Free Energy Change 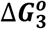 for I_2_ → Stable OC from Interactions of RNAP with Different Downstream Regions of Promoters with L- and T-Discriminators. Contributions to 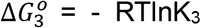 at 37 °C (in kcal/mol, see Table S4) are calculated from k_-2_ values for each promoter variant (Table S1) and k_d_ values for FL, DT+12 and DT+6 promoters (Table S2, S3). Colors and the order of promoters are the same as in previous figures.

Overall 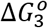 values for formation of a stable OC from I_2_ range from −6.6 kcal to −8.0 kcal for promoters with the L discriminator, and from −3.4 to −4.1 kcal for promoters with the T-discriminator (Table S4). Approximately half of the overall 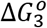 for L-discriminator variants and almost all of 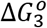 for T-discriminator variants is derived from interactions with the promoter DNA in the region from +6 upstream that are not present in I_2_. RNAP interaction free energies with this upstream region of the promoter in converting I_2_ to a stable OC are most favorable for LTL and LLL (−5.0 and −4.6 kcal, respectively). For the other six variants studied (TTL, TLL and all four T discriminator variants), interactions with this most-upstream portion of the downstream DNA contribute from −3.3 kcal to −3.8 kcal to 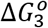.

For promoters with the L- (and to an extent T-) discriminator, Fig. 5 shows opposing trends in the contributions to 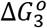 from the +7 to +12 region and the region downstream of +12. For the L-discriminator series, interactions of RNAP with the +7 to +12 region contribute most for TTL (−3 kcal) and least for LLL (−0.6 kcal), while interactions with the region downstream of +12 contribute most for LLL (−2.1 kcal) and least for TTL (−0.9 kcal). Taken together, interactions of

RNAP with the DNA downstream of +6 formed in the conversion of I_2_ to a stable OC are most favorable for TTL (−3.9 kcal) and least favorable for LLL (−2.7 kcal).

For the T-discriminator series, interactions of RNAP with the +7 to +12 region make relatively small unfavorable contributions to 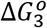. The least unfavorable contributions are observed for TTT (+0.3 kcal) and LTT (+0.1 kcal), while contributions from TLT (+0.8 kcal) and LLT (+0.6 kcal) are more unfavorable (Table S4). For this T-discriminator series, interactions with the region downstream of +12 make relatively small favorable contributions to 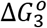. The most favorable contributions are for LLT and TLT (both −0.9 kcal) and the least favorable is for TTT (0 kcal). Taken together, interactions downstream of +6 make small net contributions to 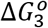 for the T-discriminator series promoters, ranging from +0.3 kcal to −0.4 kcal with uncertainties of almost 100%.

### Urea Affects Lifetimes of L- and T-Discriminator Promoters Differently

Effects of urea on equilibrium and rate constants of steps of formation and dissociation of RNAP-promoter OC, quantified as thermodynamic and kinetic *m*-values, provide unique insights into changes in water-accessible protein and nucleic acid surface area from interface formation and coupled conformational changes in those steps [17, 19, 20]. As expected from previous studies of protein folding (e.g. ref. [38, 39]) and protein-nucleic acid interactions (e.g. ref. [40]), logarithms of rate and/or equilibrium constants for association, isomerization and dissociation of RNAP from the *λ*P_R_ promoter are linear functions of urea concentration. From the slope of these plots, kinetic or thermodynamic *m*-values are determined, while the intercepts yield rate or equilibrium constant in the absence of urea.

Experimental time courses of dissociation of stable OC at different urea concentrations (0.25 – 1.5 M) are plotted in Fig. S3. Dissociation rate constants k_d_ of stable OC of L- and T-discriminator series variants obtained from analysis of these data are listed in Table S5. The corresponding OC lifetimes (τ_d_ = 1/k_d_) are plotted on a log (decade) scale vs urea concentration in Fig 6A. Lifetimes of OC for L- and T-discriminator promoters are reduced by about 25-fold and 12-fold, respectively, for each 1 M increase in [urea]. Slopes of these plots are about 30% larger for promoters with the L-discriminator than for T-discriminator promoters.

**Figure 6.**
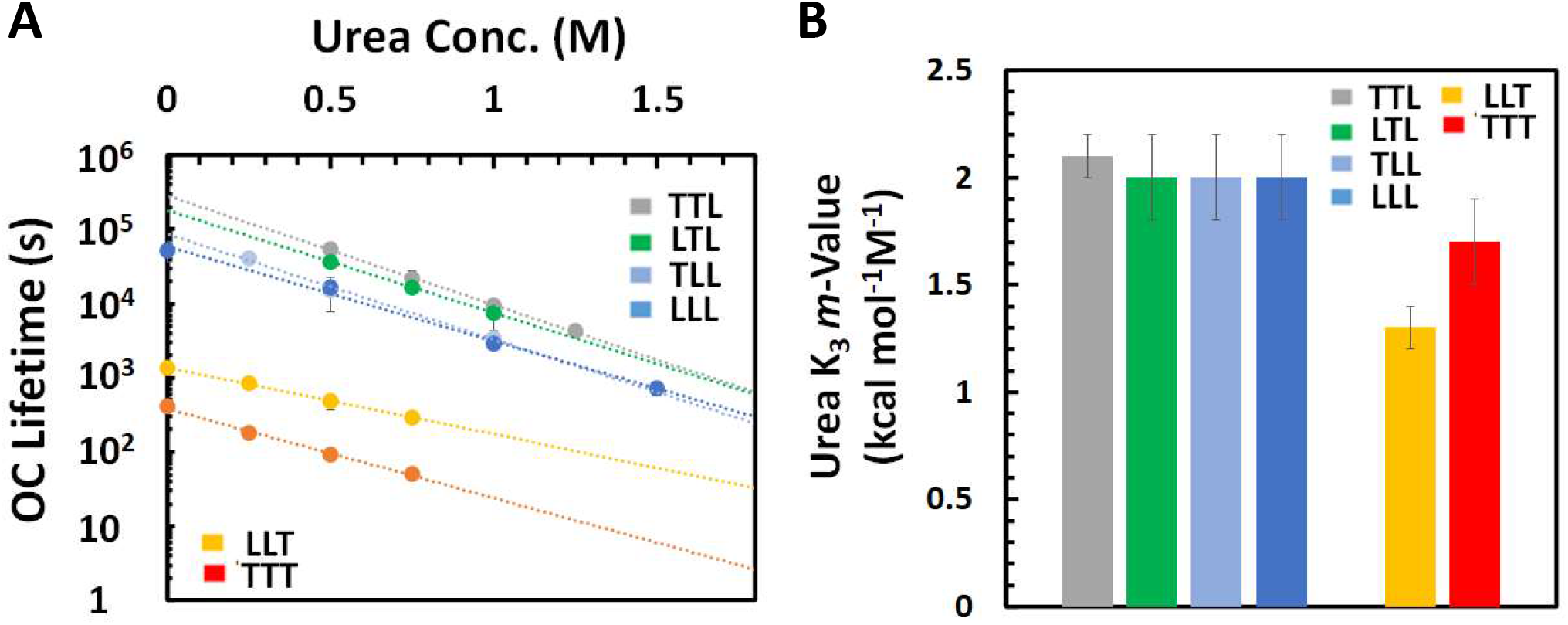
Urea Effects on OC Lifetimes for Variant Promoters with L and T Discriminators. A) Dependences of OC lifetime (in s, plotted on decade scale) on urea molarity for LLL and TTT promoters and variants. Filter binding dissociation kinetic data used to determine OC lifetimes are shown in Fig. S3. Results for LLL are from ref [17]. Extrapolation to [urea] = 0 was used to determine OC lifetimes in Fig. 2 B, C for TLL, LTL and TTL. B) Histogram of urea K_3_ *m*-values (kcal mol^-1^ M^-1^; see Table S6) quantifying the effect of urea on the equilibrium constant K_3_ for the process I_2_ → stable OC. Urea K_3_ *m*-values are obtained from the slopes of panel A and the previous finding that the rate constant k_-2_ of the DNA closing step (I_2_ → closed complexes) is independent of urea concentration [17].

Because urea does not affect the lifetime (1/k_-2_) of the I_2_ intermediate [17], the large reductions in stable OC lifetime with increasing [urea] result entirely from reductions in K_3_ for conversion of I_2_ to the stable OC with increasing [urea], and the slopes of these plots yield thermodynamic (K_3_) *m*-values. (urea K_3_ *m*-value = - RTdlnK_3_/d[urea] = RTdlnk_d_/d[urea] = - RTdlnτ_d_/d[urea] because dlnk_-2_/d[urea] = 0 [41].) These K_3_ *m*-values are compared in the bar graph of Fig. 6B (see also Table S6); K_3_ *m*-values for L-discriminator variants are about −2.1 ± 0.2 kcal mol^-1^ M^-1^ and for T-discriminator variants are about −1.5 ± 0.3 kcal mol^-1^ M^-1^.

From the previous finding at 25 °C that urea K_3_ *m*-values of the DT+12 and ΔJAW variants of the LLL promoter are about 2/3 as large as that of the FL LLL promoter (see Table S6) [20], we dissect contributions to the L-discriminator series K_3_ *m*-values as approximately −0.7 kcal mol^-1^ M^-1^ from interactions downstream of +12 and −1.4 kcal mol^-1^ M^-1^ from interactions with +12 and upstream DNA. This upstream contribution to the K_3_ *m*-value for promoters in the L-discriminator series (−1.4 kcal mol^-1^ M^-1^) is the same within uncertainty as the average K_3_ *m*-value for the T-discriminator promoters investigated (−1.5 kcal mol^-1^ M^-1^). From this equality, together with the finding that most if not all contacts made in the conversion of I_2_ to a stable OC for T-discriminator promoters are with +6 and upstream DNA, we deduce that these +6 and upstream interactions are responsible for the entire −1.4 kcal mol^-1^ M^-1^ contribution from interactions upstream of +12 to the urea K_3_ *m*-value of L-discriminator promoters. Hence, the interaction of RNAP with the promoter region between +12 and +7 in conversion of I_2_ to stable OC for the L-discriminator series is not detectably urea-dependent, and so involves no large-scale coupled folding. This dissection of the urea K_3_ *m*-value is shown in the left three columns of Table S6.

These urea K_3_ *m*-values for interactions with +6 and upstream DNA for both L- and T-discriminator series (−1.4 kcal mol^-1^ M^-1^) are very large in magnitude, as discussed below. This behavior is what is expected if there is extensive coupled folding [41], burying a large amount of previously water-accessible RNAP surface, in forming the RNAP-promoter interaction with +6 and upstream DNA in the conversion of I_2_ to a stable OC.

## Discussion

From free energy changes 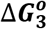 for the process of converting the initial unstable OC (I_2_) to a more stable OC (eg. I_3_, RP_O_ at λP_R_) for sets of full-length and downstream-truncated promoter variants with T- and L-discriminators, contributions to the stabilization of the open complex (I_2_ → stable OC) from interactions of downstream mobile elements (DME) of RNAP with three regions of the downstream duplex were determined (Fig. 5). In the following sections we discuss the RNAP DME (clamp, lobe, jaw) that are the most likely candidates to interact with the downstream λP_R_ duplex [3, 20, 42], and why only a subset of these elements interact with the T7A1 duplex [3]. We also discuss the differences between λP_R_ and T7A1 discriminators and upstream elements that appear to be most significant as determinants of their different downstream interactions in stable OC. These upstream determinants are quite distant from the DME-DNA interaction they affect, and we discuss the allosteric network that transmits the signal from upstream to downstream to affect the grip of RNAP on the downstream λP_R_ duplex (but not the T7A1 duplex) in I_2_ → RP_O_.

### Interactions of the +6 and Upstream Promoter DNA with β’ Switches 1 and 2 at Base of Cleft Close the β’ Clamp in Converting Unstable Open Intermediate I_2_ to a Stable OC

The largest contribution to 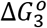 for I_2_ → stable OC is made by newly-formed interactions of RNAP with the region of promoter DNA from +6 upstream (Fig. 5, Table S4). These may include interactions with the downstream discriminator and TSS as well as the duplex from +3 to +6. These +6 and upstream interactions are of similar strength (−3 to −4 kcal) for all eight promoter variants studied. This contribution accounts for almost the entire 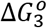 for promoters with the T-discriminator, and about half of 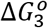 for promoters with the L-discriminator. Urea greatly destabilizes this interaction, reducing its contribution to 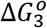 for TTT and LLT promoter complexes by ~1.5 kcal at 1 M urea (Table S6). These large effects of urea are comparable to urea effects on Δ*G*° of folding a 100-residue polypeptide chain to form a globular protein (ΔASA ≈ −8 x 10^3^ A^2^, ref. [39]), as well as to effects of urea on Δ*G*° of forming a lac repressor-lac operator complex with a 20 bp interface and coupled folding of more than 20 residues of the hinge regions (ΔASA ≈ −6 x 10^3^ A^2^, ref. [40]).

The downstream promoter duplex from +3 to +6 contacts β’ switch 1 and switch 2 (β’ residues 1304-1329 and 330-349, respectively; Fig. 7A) of the RNAP [43–45], which are located at the base of the β’ clamp (Fig. 7B) and serve as hinges for opening and closing of the clamp [28, 43, 46]. These switches each have two folded conformations, changing conformation in clamp closing [43]. Adjacent to each switch are about 20 residues of the β’ clamp that are predicted by PONDR to be unfolded (residues 1330-1350 and 309-329; [17, 20]). PONDR also predicts portions of both switches to be unfolded. While these PONDR regions are folded in holoenzyme structures and in a stable promoter complex [22, 47], they may be unfolded in I_2_ [48].

**Figure 7.**
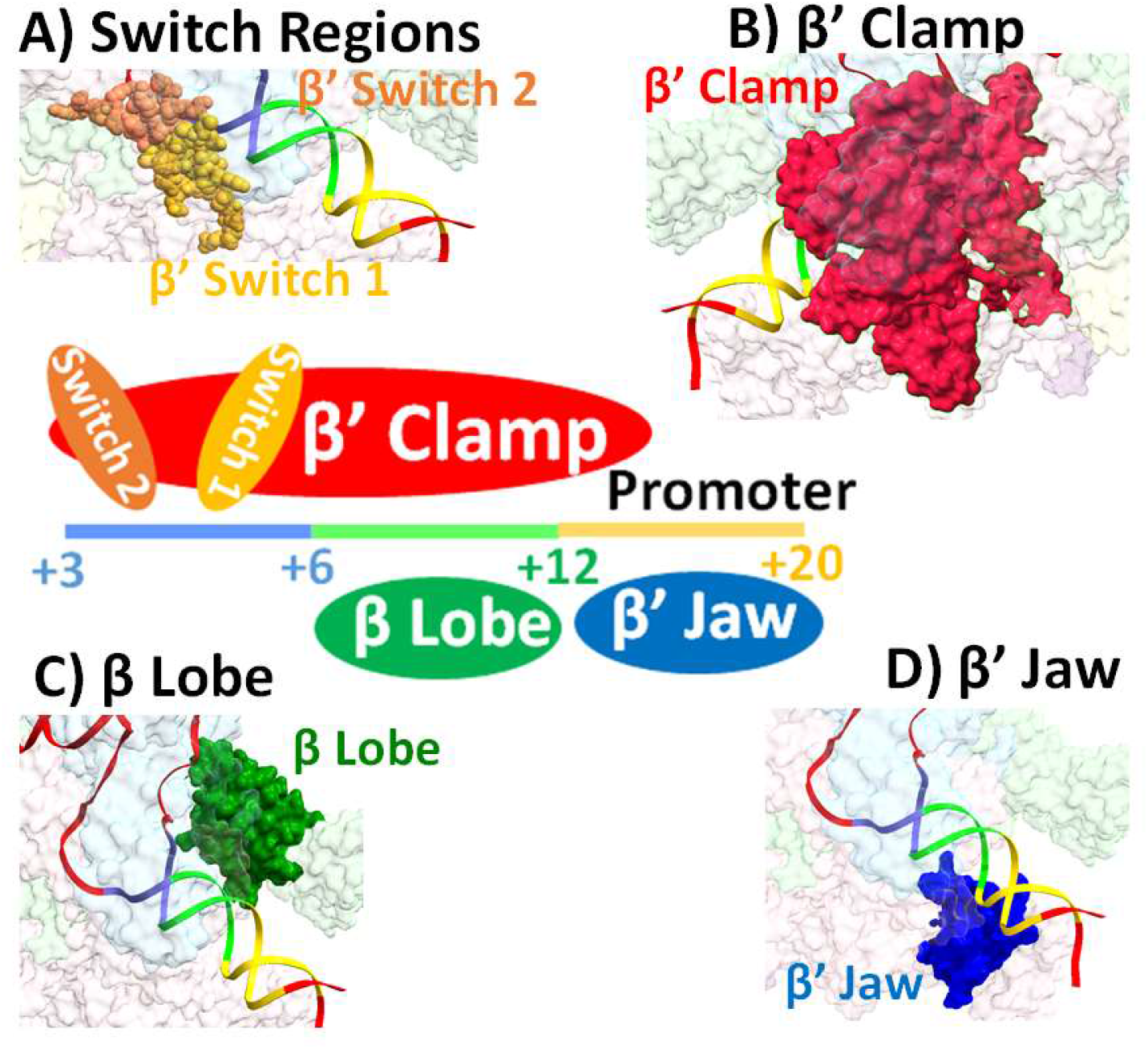
Downstream Mobile Elements (DME) Involved in OC Stabilization (I_2_ → stable OC) The schematic diagram (center of figure) shows RNAP elements and regions of the downstream promoter duplex that they interact with in OC stabilization. **Panels A, B: Mobile elements involved in clamp closing on the +3 to +6 duplex.** A) The β’ switches 1 (gold) and 2 (coral), which are hinges for the RNAP β’ clamp, interact with duplex region +3 to +6 of all L- and T-variant promoters in the conversion of I_2_ to a stable OC. B) This interaction triggers folding of adjacent elements of the β’ clamp and clamp-closing. **Panels C, D: Mobile elements involved in promoter-specific interactions downstream of +6** C) The β-lobe (green) contacts the duplex of L-discriminator (but not T-discriminator) variant promoters between +7 to +12 (light green). D) The β’-jaw (blue) contacts the duplex of L-discriminator (but not T-discriminator) variant promoters downstream of +12. (All panels are generated from PDB 7MKD [22])

For the eukaryotic pol II, direct contacts between the switch region and DNA phosphates are deduced to coordinate and perhaps mechanically couple clamp closure and DNA binding [44, 45]. Pol II crystal structures show that the downstream duplex DNA contacts and folds the switch regions, triggering the folding of the clamp domain to allow NTP binding and initiation of transcription [45]. Our results summarized above provide strong evidence for a similar mechanism in OC formation by *E. coli* RNAP and identify the mechanistic step (I_2_ → I_3_) in which it occurs.

Hence we propose that in the unstable open complex (I_2_) the β’ clamp is open and regions connecting the switches to the β’ clamp (e.g. those identified by PONDR as likely to be intrinsically disordered) are unfolded. Contact of the +3 to +6 region of the duplex with the switches causes them to change conformation and induces folding of regions connecting these switches to the rest of the β’ clamp [43], resulting in clamp-closing and stabilization of the OC by 3 - 4 kcal for both T- and L-discriminator promoters. Folding of these >40 residues, together with contacts in the interface, can explain the large-magnitude urea K_3_ *m*-values of the T-discriminator promoters, deduced to arise from this step (I_2_ → stable OC; Table S6) because these T-discriminator promoters have only limited contacts with RNAP downstream of +6 (Table S6). A possible analogy is to the lac repressor – lac operator interaction, where interaction of the flexibly-tethered DNA-binding-domains of repressor with the operator results in folding of the flexible tethers into α-helices and formation of a large interface with the core repressor [40, 49, 50].

From an analysis of changes in Cy3-to-Cy5 FRET efficiency for probes located at −100 and +14 positions of promoter DNA in the steps of λP_R_ OC formation, we previously deduced that the β’ clamp opens to admit the downstream duplex and then closes on the duplex in the most advanced CC (designed I_1L_) [6]. We proposed that clamp opening accompanies opening the downstream duplex to form the unstable I_2_ open intermediate from I_1L_. This proposal, which provides a mechanism for the use of strand binding free energy in DNA opening [6], is consistent with our current finding that the clamp is open in I_2_ and closes in the subsequent conversion of I_2_ to a stable OC.

Studies of the effect of downstream truncation of the duplex on the ability of the OC formed with N25cons and λP_R_ promoters to initiate upon NTP addition [28] revealed that DT+6 is the minimum length for initiation. Very possibly the interaction of the +3 to +6 promoter region with the switches in the conversion I_2_ → stable OC causes both clamp closing and the observed realignment of TSS bases relative to the active site (Figure S4) [13] that allows initiation. Recent transcription initiation studies with λP_R_ revealed that I_3_ or an I_3_-like complex (the stable OC at or below 20 °C) is the initiation complex, and that the stable 37 °C OC (RP_o_) cannot initiate [21].

For promoters like λP_R_, clamp closing may be coupled to DME folding and/or movements downstream, including the β lobe and β’ jaw discussed in the next section, and may also be coupled to movements in other in-cleft elements in addition to the switch regions, including the polyanionic (DNA-mimic) N-terminal region 1.1 of σ^70^ (σ_1.1_). Roles of these RNAP elements in the events that stabilize the initial OC are discussed in subsequent sections.

### Interactions of DME (β lobe, β’ jaw, downstream β’ clamp) with the Downstream Duplex (Positions +7 to +20), Absent in I_2_, Stabilize the λP_R_ OC but not T7A1 OC

Dissection of 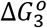 values (Fig. 5 and Table S4) reveal a significant favorable interaction of a RNAP DME with the +7 to +12 region of the downstream duplex of promoters with the L-discriminator, ranging from about −0.5 kcal to −3 kcal for the different core promoter and UP-element combinations. By contrast, interactions between RNAP DME and this region of promoters with the T-discriminator are slightly unfavorable (contributing 0 to +0.5 kcal). The β lobe (Fig. 7C) and the downstream portion of the β’ clamp (Fig. 7B) are positioned to interact with the +5 to +8 region of the downstream duplex, and both lobe and clamp can move relative to the central core of the enzyme [42, 51–54]. It is therefore likely that these elements make the favorable interactions observed with the +7 to +12 region of promoter variants with the L-discriminator. (Interactions with this region are much weaker and are unfavorable for promoters with the T-discriminator, as discussed below.) MD simulations indicate that both the β lobe and the downstream β’ clamp are highly flexible mobile elements (DME), capable of assuming multiple conformations and of functioning together to determine the path of the downstream helix [51].

The 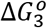 dissection in Fig. 5 also reveals a significant favorable interaction of RNAP DME with the region downstream of +12 of promoters with the L-discriminator, varying from −1 kcal to −2 kcal for the different core promoter and UP-element combinations. This interaction almost certainly involves the β’ jaw (Fig. 7D), because destabilizing contributions of similar magnitude (1 to 2 kcal) are observed when the β’ jaw and/or the duplex downstream of +12 are deleted (Fig. 4, Table S2; [3, 20]). By contrast, interactions between RNAP DME and this region of promoters with the T-discriminator are only weakly favorable (0 to −1 kcal).

In OC, the jaw is proposed to have the option of binding to the N-terminal domain of σ_1.1_, a duplex mimic, or to duplex DNA [3]. For L-discriminator promoters, it appears that the interaction of σ_1.1_ with the jaw is disrupted in the conversion of I_2_ to stable OC, allowing the jaw to bind to the region of the duplex downstream of +12 [3, 42]. For the TTT promoter, it appears that σ_1.1_ remains bound to the jaw in the conversion of I_2_ to stable OC (Fig. 8B) [3] preventing interactions of the β’ clamp (Fig. 7B), β lobe (Fig. 7C) and β’ jaw (Fig. 7D) with the downstream duplex DNA. Table S4 reveals no significant contribution to 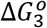 from interactions downstream of +12 (i.e. no interactions with the β’ jaw). However, the T-discriminator promoter variants (TLT, LLT) with the L core promoter exhibit some favorable interactions with the β’jaw (Fig. 8). One possible explanation of this is that the introduction of these elements of the λP_R_ promoter may lead to formation of a mixed population of stable OC and that favorable interactions with the β’ jaw are present in a minority fraction of that population.

**Figure 8.**
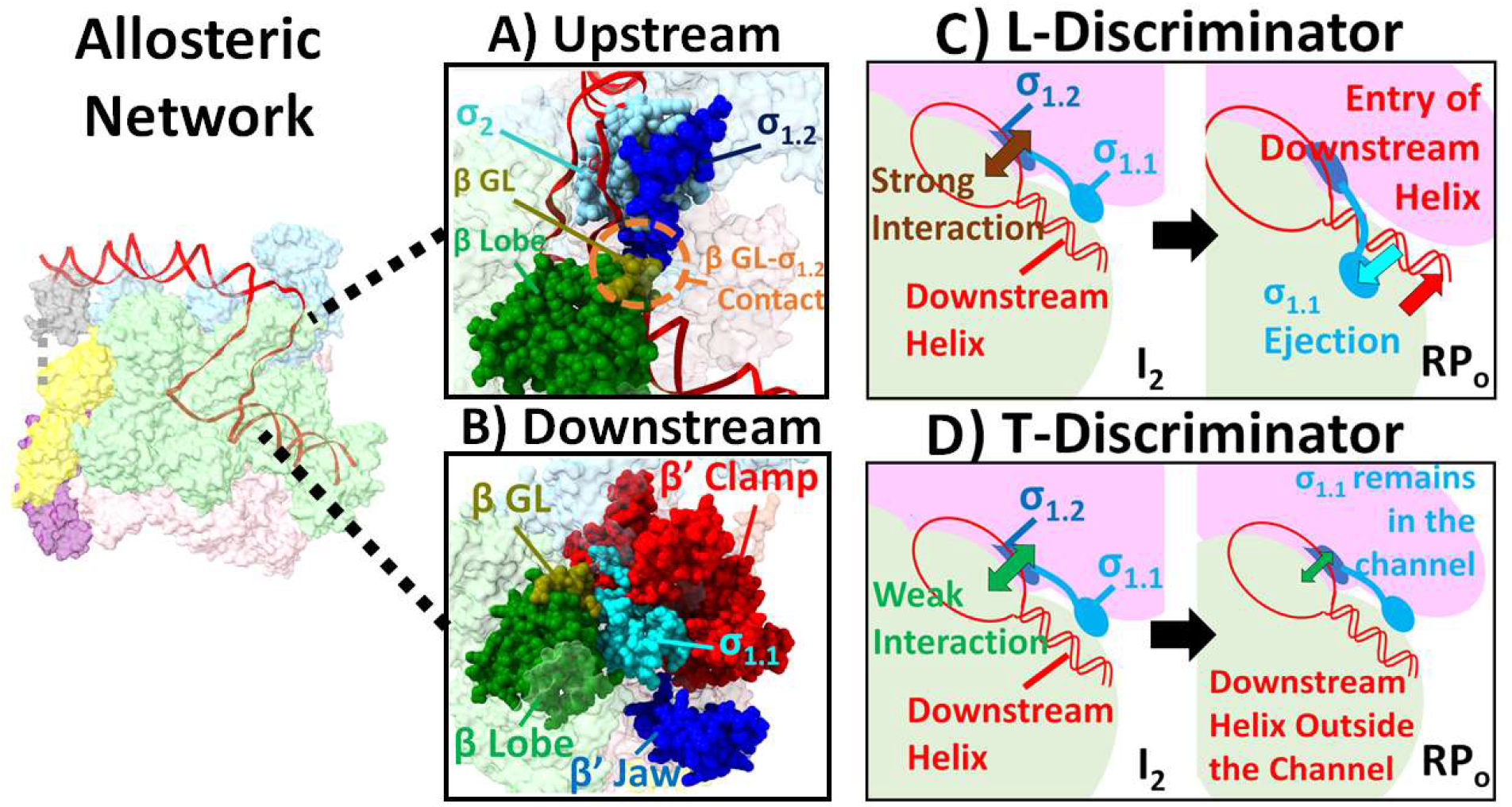
Allosteric Network that Transmits Upstream Promoter Sequence Information to RNAP Downstream Elements for Promoter-Specific OC Stabilization Downstream of +6. A) Sensing upstream promoter sequence information via interactions of σ_1.2_ and β GL with nt strand bases of the upstream discriminator. Interactions are stronger in L-discriminator variants, weaker in T-discriminator variants, and modulated by −10 region - σ_2_ interactions. Base specific interactions are transmitted downstream from σ_1.2_ to the polyanionic flexible linker and N-terminal folded domain of σ_1.1_, and from β GL to the β lobe. (Generated from PDB 7MKD [22]) B) Array of DME (β lobe, β’clamp, β’ jaw) along the downstream channel, here occupied by the N-terminal folded domain of σ_1.1_, a duplex mimic. (Generated from PDB 4YG2 [48]) C) Schematic showing that ejection of σ_1.1_ from the channel as a result of strong upstream interactions of σ_1.2_ with the L-discriminator allows the downstream duplex to enter the channel and interact with the downstream DME. D) Schematic showing that weaker upstream interactions of σ_1.2_ with the T-discriminator don’t eject the N-terminal folded domain of σ_1.1_ from the channel, preventing interactions of DME with the downstream duplex (beyond +6).

### Effects of Upstream Promoter Elements on Downstream Interactions Responsible for OC Stabilization 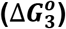

As discussed in previous sections, values of 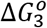 for the process of stabilizing the initial OC (I_2_ → Stable OC) at the eight full-length λP_R_ and T7A1 promoter variants investigated here differ greatly. All eight promoters exhibit similarly large contributions to 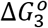 from interactions of their +3 to +6 duplex region with the β’ clamp which closes on this duplex region in this OC stabilization step (Fig. 7A and B). Differences in 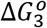 between promoters result largely from differences in interactions of RNAP DME with regions of the promoter duplex downstream of +6. These downstream interactions are regulated at a distance (allosterically) by the upstream elements of these promoters. In this section, we consider how differences in sequence or length of the discriminator, core promoter and UP element are sensed by RNAP and cause the differences in strength of downstream interactions with the β lobe and β’ clamp (+7 to +12) (Fig. 7B and C) and the β’ jaw (downstream of +12) (Fig. 7D) and possibly other DME. Input for this discussion is provided by Fig. 5 and Table S4, which dissect 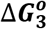 for each upstream promoter variant into contributions from interactions with the three downstream regions.

### Discriminator

The discriminator is largely responsible for the very different 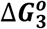 observed for the eight full-length λP_R_ and T7A1 promoter variants investigated. Table S4 shows that differences in 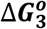 for OC stabilization between promoters with L vs. T discriminators are in the range −3 to −4 kcal (L-discriminator more favorable) irrespective of core promoter and UP element. By comparison, differences in 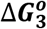 values between promoters with the same discriminator and UP element and different (L vs T) core elements span a range of approximately 1.5 kcal (from + 0.5 to −1 kcal) and differences in 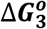 values between promoters with L vs T UP elements but the same discriminator and core elements span a range of only ~0.3 kcal (from −0.4 to −0.7 kcal; L UP element more favorable).

Differences in length and in sequence between T and L discriminators may both contribute to the large differences in their 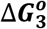 values. Considering length first, the 6 base strands of the L-discriminator (GGTTGC) appear to be the correct length to bridge between the −10 element and the TSS of an OC without scrunching or stretching [35], while pre-scrunching each strand by 1 base [34] is necessary in the case of the 7 base strands of the T-discriminator (TACAGCC). Surprisingly, no systematic effect of this magnitude (a factor of 10 change in an equilibrium constant) is observed in the equilibrium constant K_1_K_2_ (Table S1) for formation of I_2_ from RNAP and promoter DNA nor in equilibrium constants K_3,DT+6_ (Table S3) for converting I_2_ to a stable OC on promoter DNA truncated at +6. K_1_K_2_ is the same for TTT and TTL, and is larger for LLT than for LLL. K_3,DT+6_ is also the same for TTT and TTL, is two-fold larger for TLL than for TLT, four-fold larger for LLL than for LLT, and fourteen fold larger for LTL than for LTT. Only the last of these differences is of the magnitude expected from scrunching alone. Presumably the interactions of the additional nucleotide on each strand of the T discriminator with the closed clamp in the stable DT+6 OC compensate for the cost of scrunching in the other cases.

Sequence differences in the upstream part of the discriminator appear to be responsible for the very different strengths of interactions formed downstream of +6 in converting I_2_ to stable OC. As shown in Table S4, these interactions downstream of +6 contribute about half of the total 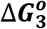 for L-discriminator variants but make no significant contribution for T-discriminator variants. The key RNAP elements responsible for recognizing the upstream discriminator sequence and transmitting its information downstream appear to be region 1.2 of σ^70^ (σ_1.2_) [29, 30] and the β gate loop (β GL) [22, 26], which are adjacent mobile elements in the cleft (Fig. 8A) [15, 22, 26].

Evidence exists for specific interactions of σ_1.2_ with the two G bases at the upstream end of the L-discriminator non-template strand [29, 30] and of the β GL with the next two bases (both T) (Fig. 8A) [26]. The T-discriminator differs in sequence at all four of these positions (TACA.. vs GGTT..) making these interactions much less favorable [26, 29, 30]. The connection of the β GL to the β lobe transmits the effect of its favorable interaction with the L discriminator downstream, positioning the β lobe and downstream β’ clamp to interact with the downstream duplex (+7 to +12) (Fig 8B).

Region σ_1.2_ is connected to the flexible tether (residues 56-98) and N-terminal three-helix bundle (residues 1–55) of region σ_1.1_ (Fig. 8B) [3, 27, 54]. These regions of σ_1.1_ are oligoanions which function as mimics of ss and ds DNA, respectively. The interaction of the L discriminator with σ_1.2_ may move or induce a conformational change in the flexible region of σ_1.1_ that repositions its N-terminal region so it no longer blocks access of the promoter duplex to the downstream cleft [3, 27, 54]. This would allow the jaw to interact with the duplex downstream of +12 [3, 5, 20, 42] (Fig. 7D). These are allosteric effects, operating over distances of 30 - 60 Å. Similar ranges of allosteric effects are observed in multi-subunit enzymes [55] and regulatory proteins [56, 57]. The lack of favorable interactions of the T-discriminator with σ_1.2_ and β GL leaves σ_1.1_ in the downstream channel, preventing entry of the duplex beyond +6, and eliminates interactions of the duplex with the lobe and clamp (Fig. 8E). These proposals expand on that made previously to explain the different lifetimes of stable T7A1 and λP_R_ OC formed with WT and deletion-variant RNAP [3, 5].

### Core Promoter and UP Element

For promoters with the L-discriminator, Fig. 5 and Table S4 reveal clear differences in contributions to OC stabilization (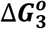 values) from interactions with the promoter duplex downstream of +6. Overall, these interactions are most stabilizing for TTL (−3.9 ±0.3 kcal), followed in rank order by LTL, TLL and LLL (which range from −3.0 to −2.7 kcal, small differences that are only marginally significant). This overall order is the result of large differences in strengths of DME interaction with the +7 to +12 regions of these promoters. Interactions with TTL are strongest (−3.0 ±0.2 kcal) and with LLL are weakest (−0.6 ±0.1 kcal). The order of interaction with DME downstream of +12 is the opposite. Interactions with LLL are strongest (−2.1 ±0.2 kcal) and with TTL and LTL are weakest (−0.9 ±0.1 kcal).

One possible explanation of these opposing patterns is that the DME, being mobile, interact with somewhat different regions of the downstream duplex in these different L-discriminator constructs, distributing their stabilizing effects differently. DME interactions with L core promoters may be focused downstream of +12, while DME interactions with T core promoters may be focused in the +7 to +12 region. Alternatively, the positions that the DME interact with on the downstream promoter DNA may be the same, but interactions with the β lobe and β’ clamp (Fig 8B) may be strengthened by the T core promoter and interactions with the β’ jaw may be strengthened by the L core promoter.

For analysis of these scenarios, a starting point is the same allosteric network as proposed to account for the discriminator effects. T and L core promoters have the same −35 element and spacer length, and differ at only one position in the −10 region. Near the downstream end, at the 5^th^ position, the L core promoter has the consensus base (A) while the T core promoter has C. A purine (G) in this position is found to interact specifically with Taq RNAP σ^A^ residues corresponding to this region of E. *coli* RNAP σ_2_ [58]. Also, purine G at this position, but not pyrimidine C, crosslinks with σ_2_ using the zero-distance crosslinker formaldehyde [59]. These findings indicate that the interaction of σ_2_ with the purine A (the L-core base) at this position is likely to be more favorable than the interaction with pyrimidine C (the T-core base). Region σ_1.2_ was proposed to stabilize a conformation of σ_2_ that is required for optimal binding of the −10 element [59]. Conversely, differences in binding of the −10 element to σ_2_ may affect the location or conformation of σ_1.2_. We therefore propose that the strength of interaction of the 5^th^ base of −10 element with σ_2_ affects the location of σ_1.2_, influencing both the β GL-σ_1.2_ contact (Fig. 8A) and the placement of σ_1.1_, with consequences for the interactions of the lobe, downstream clamp and downstream jaw with the duplex downstream of +6 (Fig. 8B).

The choice of UP element has little effect on 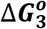 for OC stabilization. Table S4 shows that differences in overall 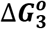 values between promoters with different UP elements and the same discriminator and core promoter elements are less than 0.7 kcal and comparable to the combined uncertainties. It may be significant that in all four comparisons 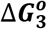 values are somewhat more favorable for the promoter with the L UP element. The largest difference in effects of L vs. T UP element is observed for promoters with the L discriminator in a comparison of interactions formed with +6 and upstream promoter DNA in conversion of I_2_ to a stable OC. Contributions from these interactions with the switches and clamp to 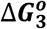 are 1 – 1.5 kcal more favorable for LLL, LTL and TLL than for TTL, for which the 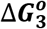 contributions are the same as for the four T-discriminator variants.

These UP-element effects appear to be on the lifetime of I_2_ before conversion to closed complexes, rather than on the subsequent stabilization of I_2_ to form an I_3_ or RP_O_-like complex. I_2_ lifetimes (1/k_-2_) are smaller for variants with the L UP element than for corresponding T UP element variants, especially for comparisons involving L-discriminator promoters. For LTL and LLL, I_2_ lifetimes are ~0.4 s, compared with ~1.2 s for TTL and ~2 s for TLL (Fig. 3 and Table S3). These UP-element effects on I_2_ lifetime are as large or larger than UP-element effects on the corresponding k_d,DT+6_ values from which K_3,DT+6_ and 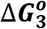 contributions from interactions with +6 and upstream are calculated (K_3,DT+6_ = k_-2_/k_d,DT+6_) (Table S3). LTL has a slightly smaller k_d,DT+6_ that the other constructs, but this is only a 2-fold difference from the mid-range value, and for LLL the k_d,DT+6_ is the mid-range value (2-fold larger than LTL, 2-fold smaller than LTT) (Table S3). Hence, the larger K_3,DT+6_ (and more negative 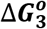 contributions) for LLL and LTL are primarily the result of larger k_-2_ values. I_2_ complexes at LLL and LTL promoters are shorter-lived that at other promoters, indicating that I_2_ is less stable at these promoters (Table S1). The more negative 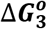 contributions from the region from +6 upstream for these promoters appear to arise because I_2_ is less stable and not because of a difference in stabilization of the final OC.

## Conclusions

From these and previous studies of the promoter and RNAP determinants of OC lifetime, we obtain answers to long-standing questions regarding why the initial open complex (I_2_) is relatively unstable and how it is subsequently stabilized at some promoters. We deduce that I_2_ is unstable because the β’ clamp is open and there are few if any interactions of RNAP DME with the downstream duplex. For λP_R_ promoter, we found previously that the clamp opens together with opening 13 bp of promoter DNA (−11 to +2, relative to the +1 TSS) in the rate-determining step in the forward mechanism [6, 60]. The lack of strong interactions with the downstream duplex at this stage (I_2_) allows it to freely rotate to dissipate the 1 1/3 turns of helical twist from opening 13 bp [60]. I_2_ is converted to a stable OC by closing the β’ clamp and, for λP_R_ but not T7A1 promoter DNA, by forming interactions of RNAP downstream mobile elements (DME; β-lobe, β’–clamp, β’-jaw) (Fig. 8B) with the duplex downstream of +6 [3, 20, 42]. Formation of these downstream interactions is directed by upstream promoter sequences via an allosteric network in which λP_R_ (but not T7A1) discriminator and −10 bases are sensed by σ_1.2_ and the β-gate loop (Fig. 8A) and this information is transmitted to σ_1.1_ and β-lobe and then to the downstream jaw and clamp (Fig. 8B) to determine OC lifetime.

## Material and methods

### Buffers

Buffers are the same as described previously [20], prepared with E-Pure deionized water and chemicals of the highest reagent grade. RNAP storage buffer contains 50% (v/v) glycerol, 0.01 M Tris (pH 7.5 at 4 °C), 0.1 M NaCl, 0.1 mM DTT and 0.1 mM Na_2_EDTA. Binding buffer (BB) contains 0.04 M Tris (pH adjusted to 8.0 at 37 °C with HCl), 0.12 M KCl, 0.01 M MgCl_2_, 1 mM DTT and 0.1 mg/ mL BSA. BB was supplemented with urea as described previously for dissociation experiments as a function of urea concentration [17]. Wash buffer (WB) for nitrocellulose filter binding assays contains 0.01 M Tris (pH 8.0 at room temperature), 0.1 M NaCl, and 0.1 mM Na_2_EDTA.

### RNAP Preparation

His-tagged but otherwise WT E. *coli* RNAP core enzyme (α_2_ββ’ω) and his-tagged △JAW RNAP (△β’ 1149-1190) core enzyme were overexpressed from E. *coli* BL21(DE3) competent cells containing either pVS10 or pIA1024 plasmid (kindly provided by Professor Irina Artsimovitch, Ohio State University) and purified using a Ni-NTA column as previously described [3]. The σ^70^ subunit was overexpressed from BL21(DE3) competent cells containing pET28 plasmid as previously described [3]. WT and △JAW RNAP holoenzymes are reconstituted by incubating the RNAP core enzyme (5 μM final) and σ^70^ (10 μM final) for 1 hr at 37 °C and stored overnight in a −20 °C freezer before use.

### Promoter DNA Preparation

To prepare 124 bp promoter DNA fragments (approximately −82 to + 42 relative to the +1 transcription start site) forward and reverse primers with length from 55 to 59 bases were designed to give promoter sequences from base −70 to +31. These forward and reverse primer sequences are complementary over a 13 base region at the 5’ end of the forward primer and 5’ end of the reverse primer. Sequences are listed in Table S6. The two primers are annealed and the single strand overhangs are filled in using One-Taq DNA polymerase (NEB). These filled-in primers are then amplified by PCR with One-Taq DNA polymerase (NEB) and two 21-base primers (HTOP and HBOT, listed in Table S2). These anneal to the 5’ end of each strand resulting in a 10 bp overlap and a 11 base overhang at the 5’ end to give further extension. The final template sequences therefore extend from −82 (for promoters with the 78 base T7A1 discriminator) or −81 to +42. Promoter fragments are purified using a PCR clean-up kit (QIAGEN) before electrophoresis on a 1.3% agarose gel (Sigma Aldrich) stained with gelstar (Lonza). Promoter bands are cut out of the gel, and subsequently purified by gel purification kit (Promega).

For downstream truncation (DT+12, DT+6) promoters, the reverse primers were shortened to the desired length, as summarized in the Downstream Truncation section of Table S6. Other procedures in the preparation of DT promoter fragments are the same as above, using the appropriate HBOT primers to obtain these lengths as listed in Table S6.

### Nitrocellulose Filter Binding Dissociation Kinetic Assay to Determine Stable OC Lifetime

Open complexes were pre-formed by incubating an excess of RNAP holoenzyme (>1.7 nM active) with <400 pM γ-^32^P-labeled promoter DNA (labeled with T4 polynucleotide kinase from NEB) in BB at 37 °C for at least 30 min. Irreversible dissociation was initiated by the addition of 50 μg/mL heparin (final concentration) or 13.4 nM unlabeled TLL promoter DNA (−82 to +42) to a sample with 36,000 cpm of promoter DNA. At selected times, 100 μL aliquots were taken and applied to 0.45 mm pre-soaked (in deionized water) nitrocellulose filters (BA85, Schleicher and Schuell, Inc., Keene, NH) on a ten-place filter manifold (Hoefer Scientific, San Francisco, CA) using a vacuum of 508-635 mm Hg. Filters were rinsed with two aliquots of 750 μL wash buffer and then immersed in scintillation cocktail (Thermo Fisher) in 7 mL disposable glass vials (Thermo Fisher) for counting on a Beckman Model LS1801 or a Hewlett Packard Tri-Carb 2100 TR scintillation counter. For each assay, background DNA retention was determined by filtering an aliquot of sample with no RNAP added. Urea dissociation kinetics experiments were performed using the same procedures as above, including urea at the desired concentration in the preincubation to form stable OC before addition of competitor, as described in ref. [17].

The dissociation rate constant k_d_ (s^-1^) was determined by fitting the background-corrected filter-retained counts from heparin-resistant (open) complexes (*cpm_t_*) as a function of time to a first order single exponential decay equation:

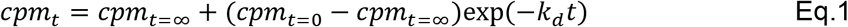

where *cpm*_*t*=0_ and *cpm*_*t*=∞_ are the values of *cpm_t_* at initial and infinite time, respectively.

### High Salt Burst Dissociation Assay to Determine the DNA Closing Rate Constant (k_-2_) and Interpret Lifetimes (1/k_d_) of Stable OC

High salt burst dissociation assays were performed on rapid mixer (Chemical-Quench-Flow model RQF-3; Kintek Co., Austin, TX) at 37 °C with one port containing pre-formed OC (RNAP > 23 nM, and promoter DNA < 600 pM) and the other port with 2.2 M KCl solution in BB as previously described [3].

From values of k_-2_ and k_d_, the equilibrium constant K_3_ for conversion of I_2_ to a stable OC and the corresponding standard free energy change 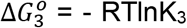 were determined from the relationship (see Figure 1 and ref. [5]).

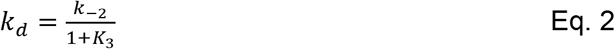

### Nitrocellulose Filter Binding Assay to Determine Overall Association Rate Constant k_a_ and the Halftime of OC Formation

To determine the kinetics of OC formation from free RNAP and promoter DNA, 10 nM of wt RNAP holoenzyme and 10 nM promoter DNA (> 50,000 cpm of γ-^32^P radio-labeled promoters supplemented with unlabeled promoter DNA of the same sequence) are mixed in BB at 37 °C at time zero. At selected time after mixing, 100 μL aliquots of samples are mixed with 20 μL heparin (50 μg/mL final), then applied to nitrocellulose filter as described above.

At 10 nM initial concentrations of RNAP and promoter DNA, concentrations of the various closed complex intermediates are calculated to be negligibly small at all times during OC formation at 37 °C, where K_1_ = (5.8 ± 2.6) x 10^6^ M^-1^ [61]. Because RNAP and promoter concentrations are equal, the kinetics of OC formation are described as by the integrated second order rate equation:

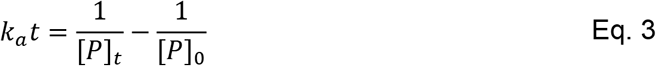

where k_a_ is the composite second order rate constant (k_a_ = K_1_k_2_; Figure 1, ref. [61]) and [P]_t_ and [P]_o_ are promoter concentrations at time t and time 0, respectively. Since [*P*]_*t*_ ≈ [*P*]_0_ − [*OC*]_*t*_, Eq. 3 rearranges to

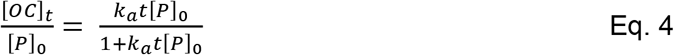

In Eq. 4, [*OC*]_*t*_/[*P*]_0_ is the fraction of promoter DNA present as OC. which is determined from the observed quantity cpmt by

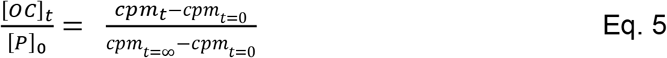

From Eq. 4 and 5, the association rate constant k_a_ (M^-1^s^-1^) is determined by fitting the background-corrected filter-retained counts from heparin-resistant (open) complexes (*cpm_t_*) as a function of time to hyperbolic Eq. 6:

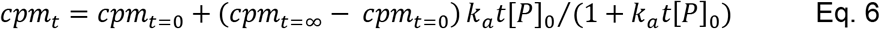

where *cpm*_*t*=0_ and *cpm*_*t*=∞_ are the values of *cpm_t_* at initial and infinite time, respectively.

## Acknowledgments

We gratefully acknowledge support of this research from NIH R35 GM 118100 and the University of Wisconsin Madison.

## Notes

### Competing Interest Statement

The authors have declared no competing interest.

