## Supplementary Information for "Roles of Clamp Closing and Allosteric Effects of Discriminator and Upstream Interactions on Downstream Elements in Stabilizing an *E. coli* RNA Polymerase-Promoter Open Complex"

Hao-Che Wang[1], Krysta Stronek[2], and M. Thomas Record, Jr.[1,2,3]

Biophysics Program[1] and Departments of Biochemistry[2] and Chemistry[3]

University of Wisconsin Madison

Madison Wisconsin 53706

**Table S1. Interactions of RNAP with T7A1 and $\lambda P_R$ Promoter Variants: Kinetic and Thermodynamic Results at 37 °C [a]**

| Construct | $k_a$ ($M^{-1}s^{-1}$) | $k_{d,FL}$ ($s^{-1}$) | $K_{obs} = k_a/k_{d,FL}$ | $k_{-2}$ ($s^{-1}$) | $K_1K_2 = k_a/k_{-2}$ | $K_{3,FL} \cong k_{-2}/k_{d,FL}$ |
|---|---|---|---|---|---|---|
| **TTT** | $(5.4 \pm 2.3) \times 10^6$ [b] | $(2.4 \pm 0.2) \times 10^{-3}$ | $(2.3 \pm 1.2) \times 10^9$ [b] | $0.6 \pm 0.3$ [c] | $(9.0 \pm 4.6) \times 10^6$ | $(2.5 \pm 1.3) \times 10^2$ |
| **LTT** | | $(1.6 \pm 0.5) \times 10^{-3}$ | | $0.8 \pm 0.4$ | | $(5.0 \pm 3.0) \times 10^2$ |
| **TLT** | | $(8.4 \pm 0.8) \times 10^{-4}$ | | $0.2 \pm 0.1$ | | $(2.4 \pm 1.2) \times 10^2$ |
| **LLT** | $(7.7 \pm 2.0) \times 10^6$ | $(7.3 \pm 0.7) \times 10^{-4}$ | $(1.1 \pm 0.3) \times 10^{10}$ | $0.6 \pm 0.4$ | $(1.3 \pm 0.9) \times 10^7$ | $(8.2 \pm 5.5) \times 10^2$ |
| **TTL** | $(1.8 \pm 1.3) \times 10^7$ | $(3.5 \pm 0.4) \times 10^{-6}$ | $(5.1 \pm 3.8) \times 10^{12}$ | $0.8 \pm 0.5$ | $(2.3 \pm 1.4) \times 10^7$ | $(2.3 \pm 1.5) \times 10^5$ |
| **LTL** | | $(5.5 \pm 0.5) \times 10^{-6}$ | | $2.4 \pm 1.0$ | | $(4.4 \pm 1.9) \times 10^5$ |
| **TLL** | | $(1.2 \pm 0.2) \times 10^{-5}$ | | $0.5 \pm 0.3$ | | $(4.2 \pm 2.6) \times 10^4$ |
| **LLL** | $(3.8 \pm 0.7) \times 10^6$ [d] | $(1.9 \pm 0.6) \times 10^{-5}$ [e] | $(2.0 \pm 0.7) \times 10^{11}$ | $2.5 \pm 1.8$ [e] | $(1.5 \pm 1.2) \times 10^6$ | $(1.3 \pm 1.0) \times 10^5$ |

[a] Symbols: FL, full-length downstream DNA; rate and equilibrium constants $\mathbf{k_a}$, $\mathbf{k_d}$, $\mathbf{k_{-2}}$, $\mathbf{K_1}$, $\mathbf{K_2}$, $\mathbf{K_3}$ **are** defined in Figure 1
[b] Ref [1] [c] Ref [2] [d] Ref [3] [e] Ref [4]

**Table S2. Dissociation Rate Constants $k_d$ for RNAP Jaw Deletion and Downstream Promoter Truncations**

| Construct | $k_{d,\Delta JAW}$ | $k_{d,DT+12}$ | $k_{d,\Delta JAW}/k_{d,FL}$ [a] | $k_{d,DT+12}/k_{d,FL}$ [b] | $k_{d,DT+6}$ | $k_{d,DT+6}/k_{d,DT+12}$ [c] |
|---|---|---|---|---|---|---|
| **TTT** | $(2.6 \pm 0.3) \times 10^{-3}$ | $(2.4 \pm 0.4) \times 10^{-3}$ | $1.1 \pm 0.2$ | $1.0 \pm 0.2$ | $(1.6 \pm 0.5) \times 10^{-3}$ | $0.7 \pm 0.2$ |
| **LTT** | $(1.3 \pm 0.2) \times 10^{-3}$ | $(3.6 \pm 0.2) \times 10^{-3}$ | $0.8 \pm 0.3$ | $2.3 \pm 0.7$ | $(2.8 \pm 0.3) \times 10^{-3}$ | $0.8 \pm 0.2$ |
| **TLT** | $(2.5 \pm 0.4) \times 10^{-3}$ | $(3.5 \pm 0.2) \times 10^{-3}$ | $3.0 \pm 0.6$ | $4.2 \pm 0.5$ | $(9.7 \pm 0.3) \times 10^{-4}$ | $0.3 \pm 0.1$ |
| **LLT** | $(1.4 \pm 0.1) \times 10^{-3}$ | $(3.1 \pm 1.5) \times 10^{-3}$ | $1.9 \pm 0.2$ | $4.3 \pm 2.1$ | $(1.3 \pm 0.1) \times 10^{-3}$ | $0.4 \pm 0.2$ |
| **TTL** | $(1.7 \pm 0.3) \times 10^{-5}$ | $(1.6 \pm 0.1) \times 10^{-5}$ | $4.9 \pm 1.0$ | $4.6 \pm 0.6$ | $(2.0 \pm 0.6) \times 10^{-3}$ | $125 \pm 40$ |
| **LTL** | $(8.0 \pm 1.7) \times 10^{-5}$ | $(2.2 \pm 0.2) \times 10^{-5}$ | $15 \pm 3.4$ | $4.0 \pm 0.5$ | $(7.0 \pm 1.0) \times 10^{-4}$ | $32 \pm 5$ |
| **TLL** | $(1.4 \pm 0.2) \times 10^{-4}$ | $(9.9 \pm 1.1) \times 10^{-5}$ | $12 \pm 2.6$ | $8.3 \pm 1.7$ | $(1.1 \pm 0.1) \times 10^{-3}$ | $11 \pm 3$ |
| **LLL** | $(5.7 \pm 2.5) \times 10^{-4}$ | $(5.5 \pm 0.7) \times 10^{-4}$ | $30 \pm 16$ | $29 \pm 10$ | $(1.4 \pm 0.2) \times 10^{-3}$ | $2.6 \pm 0.5$ |

a. $k_{d,\Delta JAW}/k_{d,DT+12} = K_{3,DT+12}/K_{3,\Delta JAW}$

b. $k_{d,DT+12}/k_{d,FL} = K_{3,FL}/K_{3,DT+12}$

c. $k_{d,DT+6}/k_{d,DT+12} \cong K_{3,DT+12}/K_{3,DT+6}$

**Table S3. Comparisons of $K_3$ Values for Downstream Promoter Truncations**

| Construct | $k_{d,FL}$ ($s^{-1}$) | $k_{-2}$ ($s^{-1}$) | $k_{-2}/k_{d,FL}$ = $K_{3,FL}$ | $k_{-2}/k_{d,DT+12}$ = $K_{3,DT+12}$ | $k_{-2}/k_{d,DT+6}$ = $K_{3,DT+6}$ |
|---|---|---|---|---|---|
| **TTT** | $(2.4 \pm 0.2) \times 10^{-3}$ | $0.6 \pm 0.3$ | $(2.5 \pm 1.3) \times 10^{2}$ | $(2.5 \pm 1.3) \times 10^{2}$ | $(3.8 \pm 2.2) \times 10^{2}$ |
| **LTT** | $(1.6 \pm 0.5) \times 10^{-3}$ | $0.8 \pm 0.4$ | $(5.0 \pm 3.0) \times 10^{2}$ | $(2.2 \pm 1.1) \times 10^{2}$ | $(2.9 \pm 1.3) \times 10^{2}$ |
| **TLT** | $(8.4 \pm 0.8) \times 10^{-4}$ | $0.2 \pm 0.1$ | $(2.4 \pm 1.2) \times 10^{2}$ | $(0.6 \pm 0.3) \times 10^{2}$ | $(2.1 \pm 1.0) \times 10^{2}$ |
| **LLT** | $(7.3 \pm 0.7) \times 10^{-4}$ | $0.6 \pm 0.4$ | $(8.2 \pm 5.5) \times 10^{2}$ | $(1.9 \pm 1.6) \times 10^{2}$ | $(4.6 \pm 3.1) \times 10^{2}$ |
| **TTL** | $(3.5 \pm 0.4) \times 10^{-6}$ | $0.8 \pm 0.5$ | $(2.3 \pm 1.5) \times 10^{5}$ | $(5.0 \pm 3.1) \times 10^{4}$ | $(4.0 \pm 2.5) \times 10^{2}$ |
| **LTL** | $(5.5 \pm 0.5) \times 10^{-6}$ | $2.4 \pm 1.0$ | $(4.4 \pm 1.9) \times 10^{5}$ | $(1.1 \pm 0.5) \times 10^{5}$ | $(3.4 \pm 1.4) \times 10^{3}$ |
| **TLL** | $(1.2 \pm 0.2) \times 10^{-5}$ | $0.5 \pm 0.3$ | $(4.2 \pm 2.6) \times 10^{4}$ | $(5.0 \pm 3.1) \times 10^{3}$ | $(4.6 \pm 2.7) \times 10^{2}$ |
| **LLL** | $(1.9 \pm 0.6) \times 10^{-5}$ | $2.5 \pm 1.8$ | $(1.3 \pm 1.0) \times 10^{5}$ | $(4.6 \pm 3.3) \times 10^{3}$ | $(1.8 \pm 1.7) \times 10^{3}$ |

**Table S4. Free Energy Contributions to $I_2$ → Stable OC from Interactions of RNAP with Different Downstream Regions (37 °C)**

| Variant | Overall (kcal) [a] | Downstream of +12 (kcal) [b] | +7 to +12 (kcal) | Upstream of +6 (kcal) |
|---|---|---|---|---|
| **TTT** | -3.4 ± 0.3 | 0 ± 0.1 | +0.3 ± 0.2 | -3.7 ± 0.4 |
| **LTT** | -3.8 ± 0.4 | -0.5 ± 0.2 | +0.1 ± 0.1 | -3.4 ± 0.3 |
| **TLT** | -3.4 ± 0.3 | -0.9 ± 0.1 | +0.8 ± 0.1 | -3.3 ± 0.3 |
| **LLT** | -4.1 ± 0.4 | -0.9 ± 0.3 | +0.6 ± 0.3 | -3.8 ± 0.4 |
| **TTL** | -7.6 ± 0.4 | -0.9 ± 0.1 | -3.0 ± 0.2 | -3.7 ± 0.4 |
| **LTL** | -8.0 ± 0.3 | -0.9 ± 0.1 | -2.1 ± 0.1 | -5.0 ± 0.3 |
| **TLL** | -6.6 ± 0.4 | -1.3 ± 0.1 | -1.5 ± 0.1 | -3.8 ± 0.4 |
| **LLL** | -7.3 ± 0.5 | -2.1 ± 0.2 | -0.6 ± 0.1 | -4.6 ± 0.5 |

[a] $\Delta G_3^o$ = - RT ln $K_3$ = RT ln ($k_{-2}/k_d$)

[b] Contribution to $\Delta G_3^o$ from interactions downstream of +12 = - RT ln ($K_{3,FL}/K_{3,DT+12}$)

[c] Contribution to $\Delta G_3^o$ from interactions between +12 and +7 = - RT ln ($K_{3,DT+12}/K_{3,DT+6}$)

[d] Contribution to $\Delta G_3^o$ from interactions upstream of +7 = - RT ln $K_{3,DT+6}$

**Table S5. Effect of Urea on OC Dissociation Rate Constant at 37 °C**

| | Dissociation Rate Constant $k_d$ ($s^{-1}$) at Urea Conc. (M) | | | | | |
|---|---|---|---|---|---|---|
| | **0 M** | **0.25 M** | **0.5 M** | **0.75 M** | **1 M** | **1.25 M** |
| **TTT** | $(2.4 \pm 0.2) \times 10^{-3}$ | $(5.6 \pm 0.4) \times 10^{-3}$ | $(1.1 \pm 0.1) \times 10^{-2}$ | $(2.0 \pm 0.1) \times 10^{-2}$ | | |
| **LLT** | $(7.3 \pm 0.5) \times 10^{-4}$ | $(1.2 \pm 0.1) \times 10^{-3}$ | $(2.1 \pm 0.4) \times 10^{-3}$ | $(3.4 \pm 0.2) \times 10^{-3}$ | | |
| **TTL** | | | $(1.9 \pm 0.1) \times 10^{-5}$ | $(4.6 \pm 1.2) \times 10^{-5}$ | $(1.1 \pm 0.2) \times 10^{-4}$ | $2.4 \times 10^{-4}$ |
| **LTL** | | | $(2.7 \pm 0.5) \times 10^{-5}$ | $(6.0 \pm 0.1) \times 10^{-5}$ | $(1.4 \pm 0.5) \times 10^{-4}$ | |
| **TLL** | | $(2.4 \pm 0.1) \times 10^{-5}$ | $(6.7 \pm 3.2) \times 10^{-5}$ | | $(2.9 \pm 0.5) \times 10^{-4}$ | |

**Table S6. Urea Effects on $k_d$ and on $K_3$ for the OC Stabilization Step $I_2$ → Stable OC at 37 °C and 25 °C**

| Construct | $d\ln k_d/d[urea]$ [a] ($M^{-1}$) | Urea $K_3$ *m*-Value[a] (kcal $mol^{-1} M^{-1}$) | Proposed Contributions to Urea $K_3$ *m*-Value from: Downstream of +12 (kcal $mol^{-1}$ $M^{-1}$) | From +7 to +12 (kcal $mol^{-1}$ $M^{-1}$) | Upstream of +6 (kcal $mol^{-1}$ $M^{-1}$) |
|---|---|---|---|---|---|
| **TTT [b]** | 2.8 ± 0.3 | 1.7 ± 0.2 | ~ 0 [b] | ~ 0 | ~ 1.5 [b] |
| **LLT [b]** | 2.1 ± 0.2 | 1.3 ± 0.1 | ~ 0 [b] | ~ 0 | ~ 1.5 [b] |
| **TTL [b]** | 3.4 ± 0.1 | 2.1 ± 0.1 | ~ 0.7 [b] | ~ 0 | ~ 1.4 [b] |
| **LTL [b]** | 3.2 ± 0.3 | 2.0 ± 0.2 | ~ 0.7 [b] | ~ 0 | ~ 1.4 [b] |
| **TLL [b]** | 3.3 ± 0.3 | 2.0 ± 0.2 | ~ 0.7 [b] | ~ 0 | ~ 1.4 [b] |
| **LLL [c]** | 3.3 ± 0.2 | 2.0 ± 0.2 | ~ 0.7 [b] | ~ 0 | ~ 1.4 [b] |
| **LLL[d]** | 3.1 ± 0.1 | 1.8 ± 0.1 | ~ 0.6 [d] | ~ 0 | ~ 1.2 [d] |
| **LLL DT+12 [d]** | 1.9 ± 0.2 | 1.1 ± 0.1 | ~ 0 | ~ 0 | ~ 1.2 [d] |
| **LLL-ΔJAW [d]** | 2.1 ± 0.1 | 1.2 ± 0.1 | ~ 0 | ~ 0 | ~ 1.2 [d] |
| **LLL DT12-ΔJAW [d]** | 2.1 ± 0.1 | 1.2 ± 0.1 | ~ 0 | ~ 0 | ~ 1.2 [d] |

a. urea $K_3$ *m*-value = - $RTd\ln K_3/d[urea]$ = $RTd\ln k_d/d[urea]$ because $d\ln k_{-2}/d[urea] = 0$ [4]

b. 37 °C (this study)

c. 37 °C [4]; $K_3 = k_3/k_{-3}$ (see Figure 1); for LLL, - $RTd\ln k_3/d[urea]$) = 1.4 ± 0.1 and - $RTd\ln k_{-3}/d[urea]$) = 0.7 ± 0.1

d. 25 °C [5]

### Table S7. Primer and Template Sequences

Primer Sequences

| | |
|---|---|
| LL-x forward | 5'-CCA CGA ATT CGG ATA AAT ATC TAA CAC CGT GCG TGT TGA CTA TTT TAC CTC TGG CGG TG-3' |
| TT-x forward | 5'-CCA CGA ATT CAA TTT AAA ATT TAT CAA AAA GAG TAT TGA CTT AAA GTC TAA CCT ATA G-3' |
| TL-x forward | 5'-CCA CGA ATT CAA TTT AAA ATT TAT CAA AAA GAG TA T TGA CTA TTT TAC CTC TGG CGG TG-3' |
| LT-x forward | 5'-CCA CGA ATT CGG ATA AAT ATC TAA CAC CGT GCG TG T TGA CTT AAA GTC TAA CCT ATA G-3 |
| x-LL reverse | 5'- ACA AAA GCT TCA TAC AAC CTC CTT AGT ACA TGC AAC CAT TAT CAC CGC CAG AGG T-3' |
| x-TT reverse | 5'-ACA AAA GCT TCA TAC AAC CTC CTT AGT ACA TGG CTG TAA GTA TCC TAT AGG TTA GAC-3' |
| x-LT reverse | 5'- ACA AAA GCT TCA TAC AAC CTC CTT AGT ACA TGG CTG TAA TTA TCA CCG CCA GAG GT-3' |
| x-TL reverse | 5'- ACA AAA GCT TCA TAC AAC CTC CTT AGT ACA TGC AAC CAT TAT CAC CGC CAG AGG T-3' |
| TT-x UT-65 forward | 5'-A ATT CAA TTT AAA ATT TAT CAA AAA GAG TAT TGA CTT AAA GTC TAA CCT ATA G-3' |
| LT-x UT-65 forward | 5'-A ATT CGG ATA AAT ATC TAA CAC CGT GCG TGT TGA CTT AAA GTC TAA CCT ATA G-3' |

Downstream Truncation

| | |
|---|---|
| x-TT_+12_Reverse | 5'- TCC TTA GTA CAT GGC TGT AAG TAT CCT ATA GGT TAG AC-3' |
| x-LT_+12_Reverse | 5'-TCC TTA GTA CAT GGC TGT AAT TAT CAC CGC CAG AGG T-3' |
| x-LL_+12_Reverse | 5'-TCC TTA GTA CAT GCA ACC ATT ATC ACC GCC AGA GGT-3' |
| x-TL_+12_Reverse | 5'- TCC TTA GTA CAT GCA ACC AGT ATC CTA TAG GTT AGA C-3' |
| x-TT_+6_Reverse | 5'- GTA CAT GGC TGT AAG TAT CCT ATA GGT TAG AC-3' |
| x-LT_+6_Reverse | 5'-GTA CAT GGC TGT AAT TAT CAC CGC CAG AGG T-3' |
| x-LL_+6_Reverse | 5'-GTA CAT GCA ACC ATT ATC ACC GCC AGA GGT-3' |
| x-TL_+6_Reverse | 5'-GTA CAT GCA ACC AGT ATC CTA TAG GTT AGA C-3' |

HTOP and HBOT

| | |
|---|---|
| HTOP | 5'-CCA GCA TTC CTC CAC GAA TTC-3' |
| HBOT | 5'-CAC CTG CAC CGA CAA AAG CTT-3' |
| HBOT_+12_T-DISC | 5'-TCC TTA GTA CAT GGC-3' |
| HBOT_+12_L-DISC | 5'-TCC TTA GTA CAT GCA-3' |
| HBOT_+6_x-TT | 5'- GTA CAT GGC TGT AAG-3' |
| HBOT_+6_ x-LT | 5'- GTA CAT GGC TGT AAT-3' |
| HBOT_+6_x-LL | 5'- GTA CAT GCA ACC ATT-3' |
| HBOT_+6_x-TL | 5'- GTA CAT GCA ACC AGT-3' |

For DT+12 constructs, the HBOT_+12_T-DISC is for promoters with T discriminator, and HBOT_+12_L-DISC is for promoters with L discriminator. For DT+6 constructs, the HBOT_+6_x-ab primers are applied, where x can be either T or L UP-element, and a and b are either L or T in the tables, indicating the product promoters' core-region and discriminator sequences, respectively.

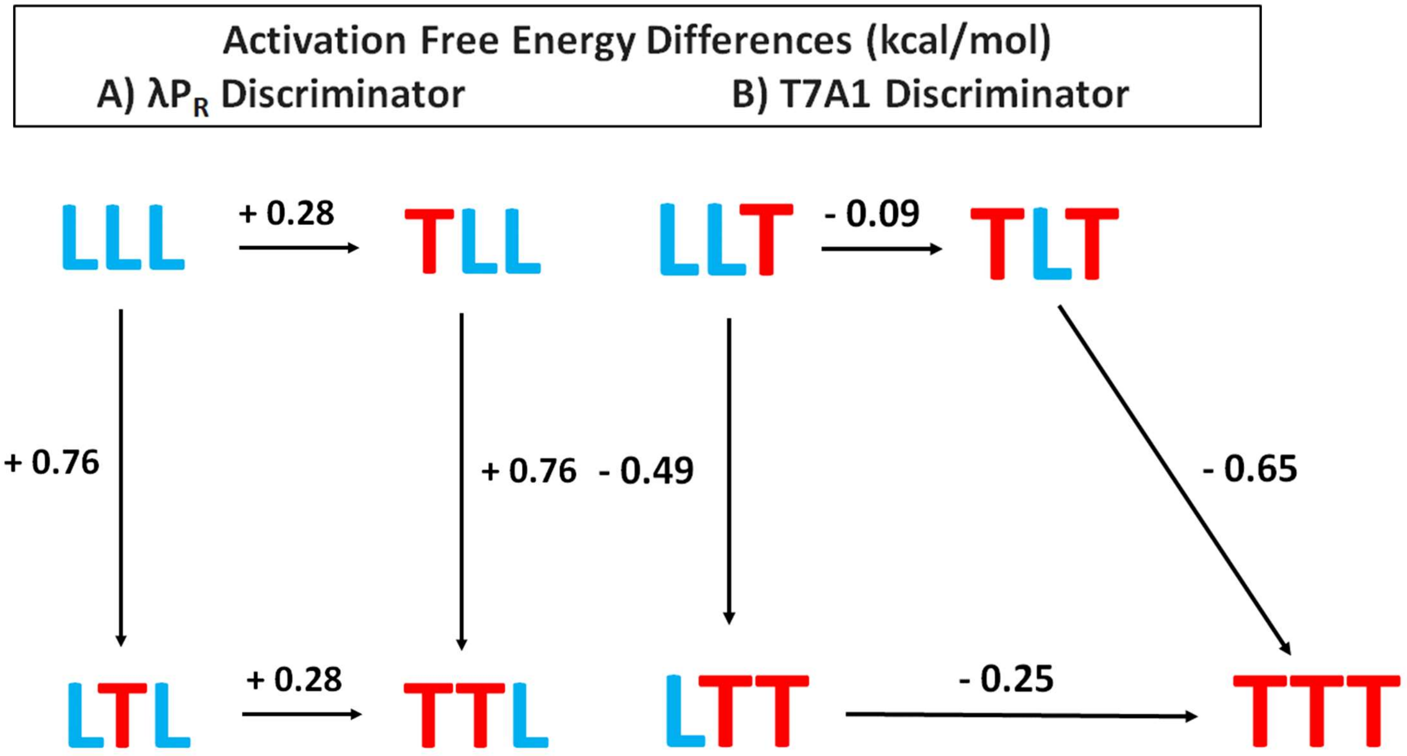

**Figure S1. Quasi-thermodynamic Cycle Diagrams of Activation Free Energy Differences $\Delta\Delta G_i^{o\ddagger}$ between Promoters with L and T Discriminators**

Activation free energy differences $\Delta\Delta G_i^{o\ddagger}$ = RTΔln$k_d$ (kcal/mol) are listed on the figure, and indicated by arrow length. A) L-discriminator OC, with LLL at upper left and TTL at lower right. B) T-discriminator OC, with LLT (the corresponding promoter in the T discriminator series to LLL) at upper left and TTT at lower right.

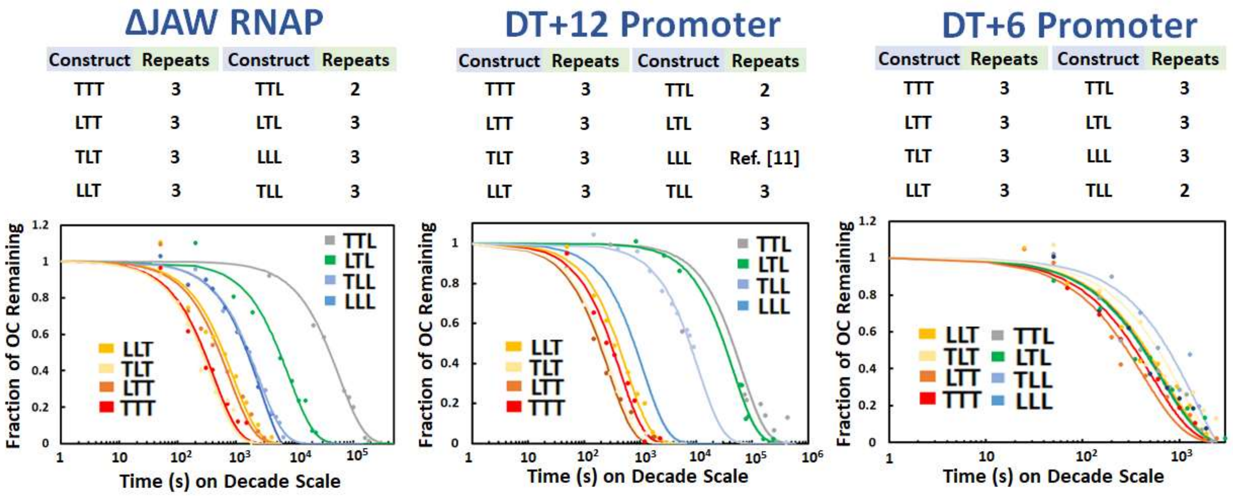

**Figure S2. Effects of Deletion of the RNAP Downstream Jaw and of Downstream Truncation of the Promoter on the Kinetics of Dissociation of Stable OC at 37 °C**

Panels list numbers of experiments for each promoter variant and show a representative OC dissociation kinetics experiment (plotted vs. time on decade scale) for each variant. Left: Dissociation of OC formed by full-length promoter DNA and the jaw-deletion (ΔJAW) RNAP variant. Center and Right: Dissociation of OC formed by downstream-truncated (DT+12, DT+6) promoter DNA and wild-type RNAP.

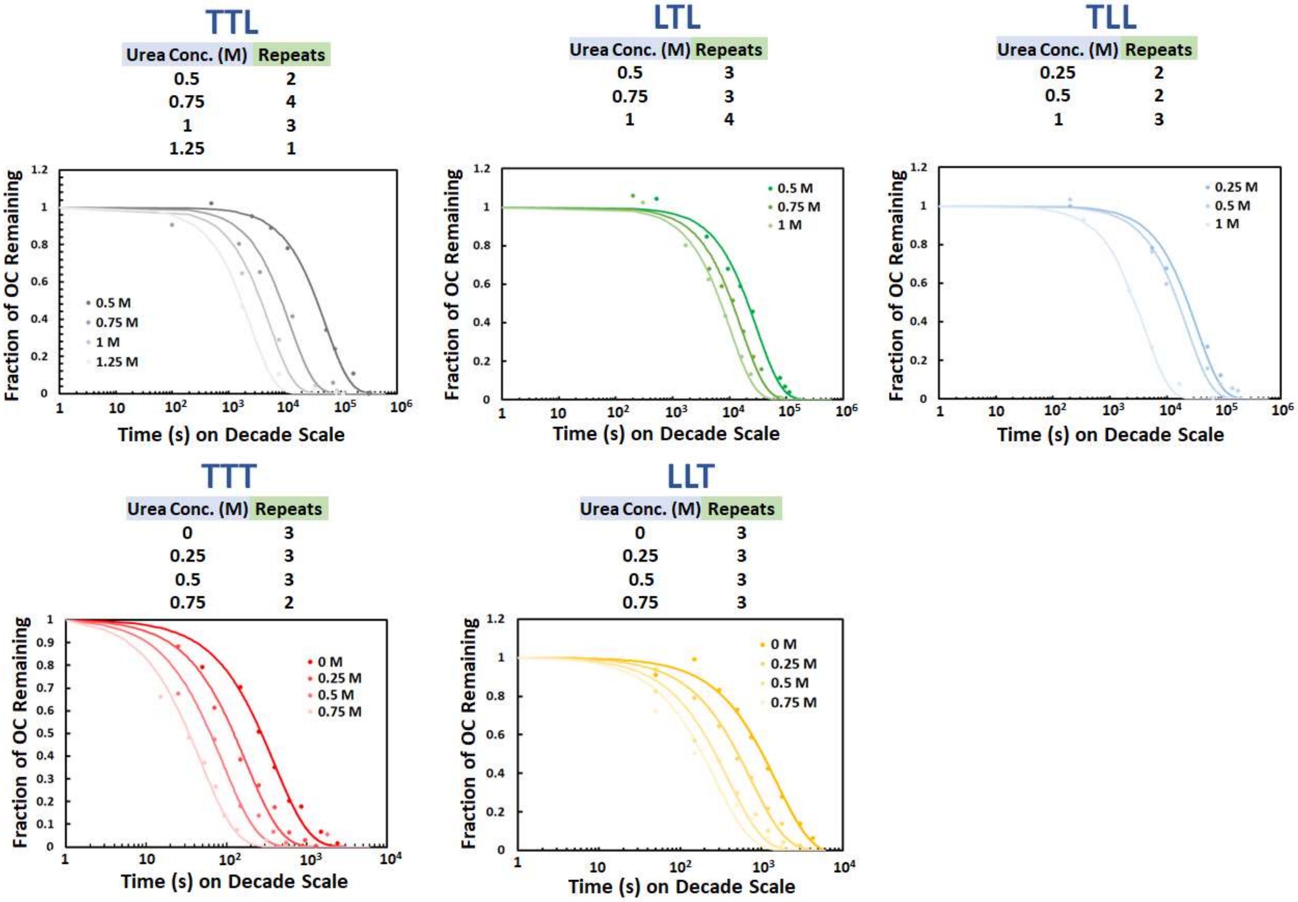

**Figure S3. Effects of Urea on the Kinetics of Dissociation of Stable OC at 37 °C**

Panels list numbers of experiments at each urea concentration and show results of representative OC dissociation kinetics experiments for each promoter variant and urea concentration investigated. Top panels: L-discriminator promoters (TTL, LTL, TLL; results for LLL were reported previously [4]); bottom panels: T-discriminator promoters (TTT and LLT).

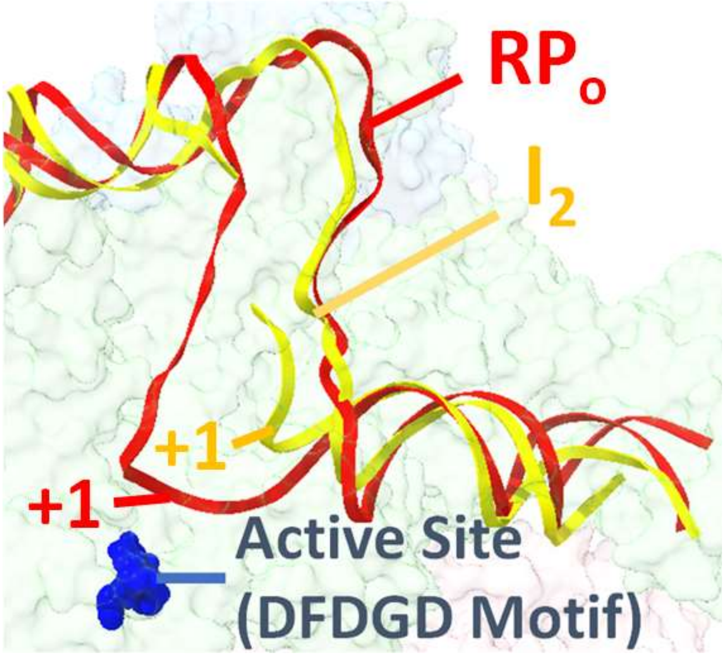

**Figure S4. Movement of the +1 TSS to the RNAP Active Site In the Conversion $I_2$ → $RP_o$.**

Shift in position of the template strand transcription start site (TSS, position +1) to the RNAP active site (blue DFDGD motif) in the conversion of the $I_2$ open intermediate (yellow DNA strands; obtained from PDB: 6EE8 [6]) to the stable OC ($\lambda P_R$ $RP_o$; red DNA strands; obtained from PDB:7MKD [7]).
